# Orfamide-A-mediated bacterial-algal interactions involve specific Ca^2+^ signalling pathways

**DOI:** 10.1101/2021.07.24.453618

**Authors:** Yu Hou, Yuko Bando, David Carrasco Flores, Vivien Hotter, Bastian Schiweck, Hans-Dieter Arndt, Maria Mittag

## Abstract

The antagonistic bacterium *Pseudomonas protegens* secretes the cyclic lipopeptide orfamide A, which triggers a Ca^2+^ signal, causing the deflagellation of the green microalga *Chlamydomonas reinhardtii*. By investigating targeted synthetic orfamide A variants and inhibitors, we found that at least two Ca^2+^-signalling pathways and TRP channels are involved in this response.

## Main text

Bacterial cyclic lipopeptides (CLiPs) are produced by several bacterial genera such as *Bacillus*, *Streptomyces* or *Pseudomonas.*^1–3^ These natural products display a wide array of biological activities. Foremost, many of these lipidated cyclopeptides, such as daptomycin,^4^ interact with lipid components of membranes and destabilize them.^1,2,5^

The CLiP orfamide A (**1**, Fig. 1) is secreted by *Pseudomonas protegens* Pf-5, a bacterium antagonistic to the green alga *Chlamydomonas reinhardtii*.^6^ *C. reinhardtii* is a genetically well-characterized model alga that bears two flagella, also known as cilia, for its movement.^7^ *P. protegens* encircles the algal cells, presumably resulting in a higher concentration of orfamide A in the microenvironment of *C. reinhardtii*.^6^ Exposure of the algal cells to orfamide A causes immediate cytosolic Ca^2+^ elevation, based on a Ca^2+^ influx for which external calcium is imported into the cell, resulting in rapid deflagellation of the algal cells (within one minute).^6^ This prevents the escape of the algae from the bacterial attack. Former data indicate that the algal cell membrane integrity is not affected. Experiments with the non-specific Ca^2+^ channel inhibitor La^3+^ suggest selective Ca^2+^ channel activity for orfamide A. This is further corroborated by the specificity of orfamide A in immobilizing Chlorophyceae algae, but not select species from the Pedinomonaceae or Euglenophytes.^6^ Ca^2+^ regulates different response signals in *C. reinhardtii*, including exposure to organic acids, osmotic stress, and light-signalling pathways.^6,8–10^ Each Ca^2+^ signalling cascade differs, and not all of them cause deflagellation. For example, salt stress- and light-induced Ca^2+^ signalling pathways do not cause shedding of the flagella, whereas an abrupt decrease of pH or the action of orfamide A rapidly deflagellate the cells. These findings imply that the algal Ca^2+^ signatures must have particular signalling mechanisms^8^ and may involve different Ca^2+^ signalling pathways.

**Fig. 1.**
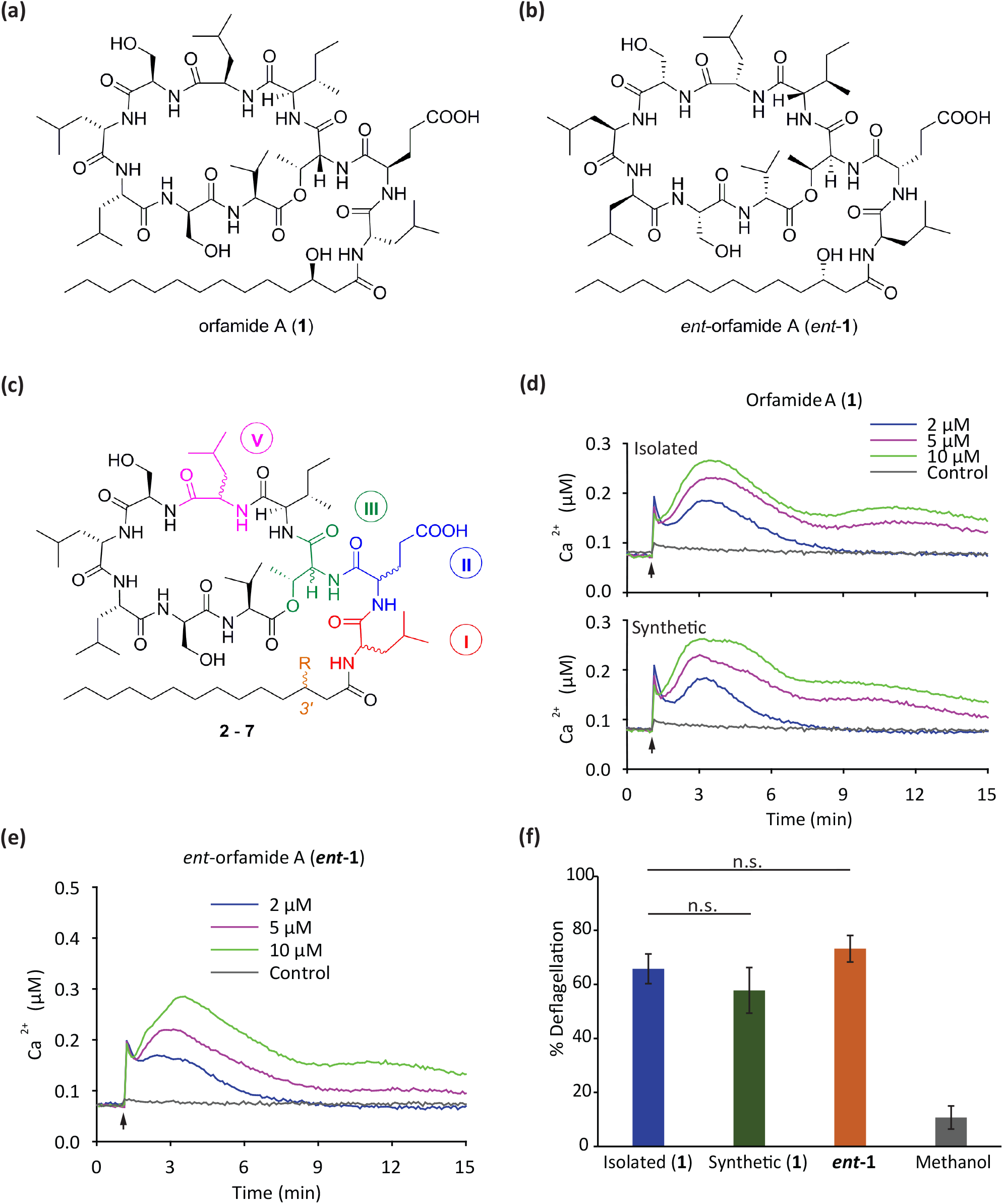
Chemical structures of synthesized (a-c) orfamide A variants as well as their biological activities (d-f). (a) Synthesized orfamide A equivalent to the isolated commercial compound. (b) Enantiomer of orfamide A. Synthesized orfamide A variants (see Table 1). (d, e) Using a *C. reinhardtii* aequorin reporter line (see Supporting Information for details), the effects of isolated and synthetic orfamide A (d, upper and lower part, respectively) as well as of the enantiomer (e) on Ca^2+^ signalling were analysed. Applied compound concentrations are indicated. Orfamide A was dissolved in methanol and further diluted in TAP medium that was added to cells at the indicated final concentrations. As control, methanol proportional to 10 μM of orfamide A and enantiomer was used. Each line represents the mean of three independent biological replicates, and each biological replicate includes three technical replicates. (f) Deflagellation rates of wild-type algal cells treated with non-saturating concentrations (5 μM) of isolated or synthetic orfamide A, its synthesized enantiomer and the solvent methanol as control. More than 100 cells were counted for each sample. Each column represents the mean of at least three independent biological replicates. Error bars represent standard deviations; Student’s t test was performed; n.s., not significant. For further details see the Supporting Information.

Chemical synthesis of some CLiPs related to orfamide A has been achieved and has facilitated their study.^11,12^ By total synthesis and validation of the biological activity of orfamide A, we^13^ and others^14^ have recently found that its originally assigned structure^15^ was stereochemically misassigned. In the correct assignment, Leu-5 features D- and the 3’-OH-group at the fatty acid chain (*R*)-configuration (Fig. 1a, Table 1).^13^ The Ca^2+^ signature induced by the correctly assigned compound of this configuration is identical to the commercially available substance, which is isolated from *P. protegens* Pf-5. Notably, this compound triggers a bimodal Ca^2+^ release in *C. reinhardtii* after exposure (Fig. 1d). In contrast, synthetic material featuring the originally reported stereochemistry shows a different induction profile, especially lacking the second Ca^2+^-release peak (Fig. 2a).

**Table 1.**
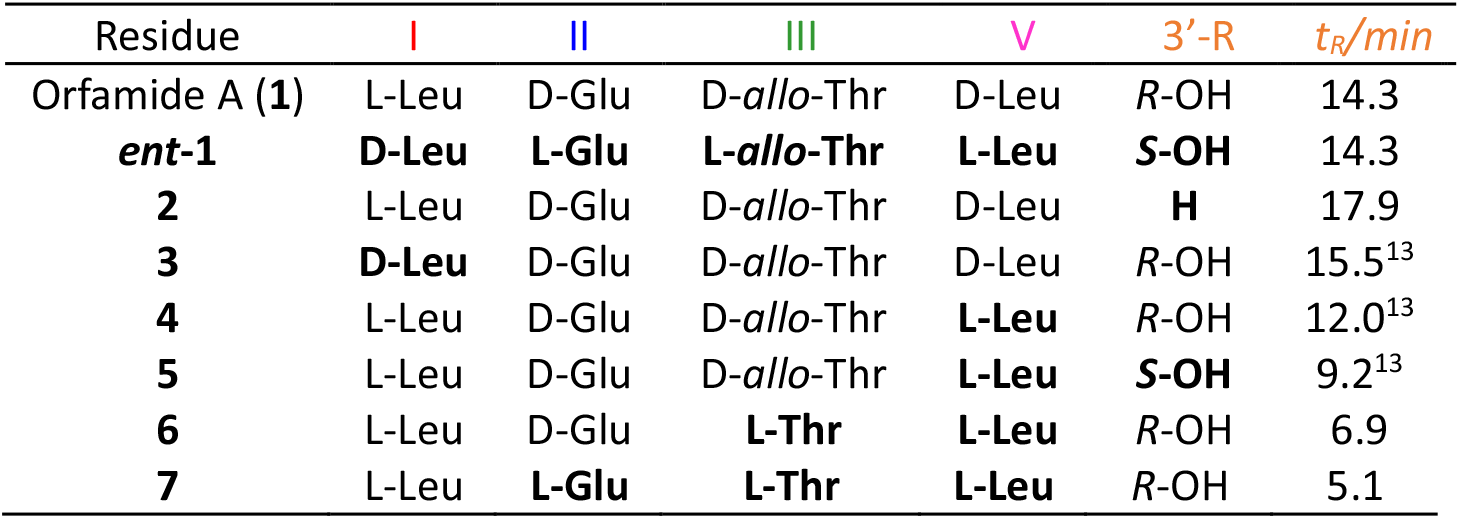
Variations of orfamide A derivatives. Changes are highlighted in bold. For HPLC conditions see Supporting Information.

| Residue | I | II | III | V | 3'-R | *t<sub>R</sub>*/min |
| --- | --- | --- | --- | --- | --- | --- |
| Orfamide A (**1**) | L-Leu | D-Glu | D-*allo*-Thr | D-Leu | *R*-OH | 14.3 |
| **ent-1** | **D-Leu** | **L-Glu** | **L-*allo*-Thr** | **L-Leu** | **S-OH** | 14.3 |
| **2** | L-Leu | D-Glu | D-*allo*-Thr | D-Leu | **H** | 17.9 |
| **3** | **D-Leu** | D-Glu | D-*allo*-Thr | D-Leu | *R*-OH | 15.5<sup>13</sup> |
| **4** | L-Leu | D-Glu | D-*allo*-Thr | **L-Leu** | *R*-OH | 12.0<sup>13</sup> |
| **5** | L-Leu | D-Glu | D-*allo*-Thr | **L-Leu** | **S-OH** | 9.2<sup>13</sup> |
| **6** | L-Leu | D-Glu | **L-Thr** | **L-Leu** | *R*-OH | 6.9 |
| **7** | L-Leu | **L-Glu** | **L-Thr** | **L-Leu** | *R*-OH | 5.1 |

**Fig. 2.**
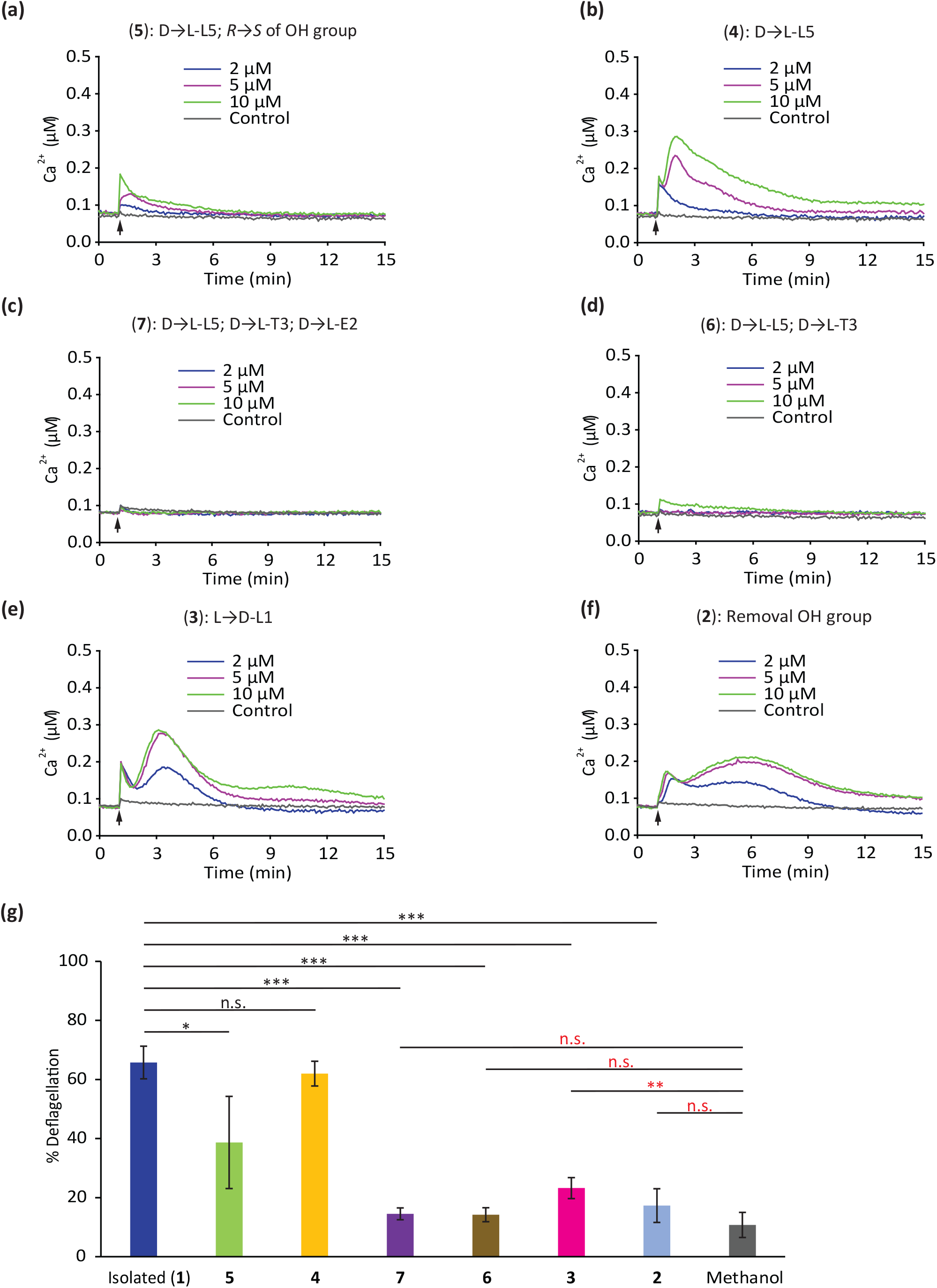
Effects of orfamide A variants on Ca^2+^ signalling (a-f) and deflagellation (g) suggest more than one Ca^2+^ signalling pathway. See legend of Figure 1 for details. Asterisks indicate significant differences as calculated by Student’s t test (n.s., not significant; *, P<0.05; **, P<0.01; and ***, P<0.001).

We have synthesized more variants of orfamide A to study their impact on biological activity upon exposure, including changes in cytosolic Ca^2+^, as well as the deflagellation phenotype. The time-dependent motility of algal cells upon compound treatment was also subjected to video analysis, as detailed in the Supporting Information. Compounds were studied at a final concentration of 10 μM, known to deflagellate >90% of the algal cells within one minute.^6^ As positive control, commercial orfamide A was used; an equal volume of pure MeOH served as negative control.

All analogues of orfamide A were obtained by total synthesis and fully characterised (see Supporting Information for details). Chemical structures of all tested compounds are compiled in Fig. 1a–c and Table 1. Importantly, treatment with synthetic and isolated (commercial) orfamide A were found indistinguishable with respect to cell motility (videos 1 and 2). It should be noted that the first sharp Ca^2+^ peak matches with the deflagellation event (cell immotility) as evident from synchronizing the Ca^2+^ response to it (videos 1 and 2). The bioactivity of the enantiomer of orfamide A (for structure see Fig. 1b) was analysed next. Interestingly, *ent*-**1** triggered a Ca^2+^ signature and deflagellation rate quite similar to **1** (Fig. 1e, f). The motility check (video 3) showed that the enantiomer may even immobilize the algal cells quicker than orfamide A (videos 1 and 2). These data indicate that for inducing the selective Ca^2+^-signal, (i) a molecular interaction with a receptor-like target may only be indirectly involved, or (ii) that only certain regions of the orfamide A molecule trigger the response. Noteworthy, the enantiomer of the cyclic lipopepsipeptide pseudodesmin A also retains activity^11^ while enantiomers of daptomycin lose activity.^12^ These data suggest that different CLiPs vary in their mechanism of biological activities.

We then analysed the compound with the formerly published incorrect configuration (**5**: L-Leu5, 3’-S-OH; Table 1). As mentioned above, the Ca^2+^ signature was different in this case (Fig. 2a). Furthermore, the deflagellation rate was significantly reduced compared to the natural compound **1** (Fig. 2g). Compound **5** did not cause significant immobilization when the cells were treated with 10 μM of the substance for up to 5 min (video 4). In a next step, we performed the assays with compound **4** which has only a L-configured Leu5 residue in the macrocycle. The data showed that this sole difference is not influencing the biological activity, as the Ca^2+^ signature resembles that of **1** (Fig. 2b), and the deflagellation rate remains similar (Fig. 2g). These results pointed - like the enantiomer (possibility ii) - to the significance of the linear segment of **1**, which was therefore studied in more detail.

Three amino acids outside of the macrocycle precede the *N*-terminal fatty acid tail: L-Leu1, D-Glu2 and D-*allo*-Thr3. Beyond the D**→**L change of Leu5, Glu2 (D**→**L) and Thr3 (D-*allo***→**L) were implemented. This triple variant **7** did not elicit a Ca^2+^ signal anymore (Fig. 2c), and deflagellation rate as well as the motility were the same as with the negative control (Fig. 2g, video 5). Interestingly, it should be noted that the variant **7** represents the most polar molecule in the series, as delineated from analytical RP-HPLC data (Table 1).

In variant **6**, only Leu5 and Thr3 were changed D**→**L (Fig. 1c, Table 1). Here, the Ca^2+^ signature was also strongly reduced (Fig. 2d). Only with 10 μM of compound, a small first Ca^2+^ peak became detectable while the second peak was fully absent. A strongly reduced deflagellation rate was observed (Fig. 2g), highlighting the importance of the Thr3 configuration.

In all these cases, the Ca^2+^ signature change correlated with the obtained deflagellation rates. This correspondence was lost when further modifications were introduced (Fig. 2e, f, g). The configurational change of Leu1 present in the linear region (L**→**D), resulted in the more hydrophobic compound **3** that gave a strong Ca^2+^ signal with bimodal shape, but showed a strongly reduced deflagellation rate (Fig. 2e, g). This was also the case when the OH group of the fatty acid tail was removed (→**2**, Fig. 2f, g). Compound **2** showed a heavily delayed response, only beginning to affect cell motility after 5 min (video 6). It could be that changes in the physicochemical properties of the compounds can modulate their potential for membrane-associated activity.^3^ Indeed, apparently minuscule changes in compound stereochemistry may profoundly influence the overall compound polarity, as indicated by C18 RP HPLC elution properties. These data suggest conformational properties and aggregation seem to be tightly involved.^11^ Furthermore, the enantiomer ***ent*-1** features full activity, rendering a direct interaction of the whole compound with a receptor-like target not very likely.

However, if general lipophilicity is analysed, no clear correlation with compound activity could be seen (Table 1). These data suggest either that the linear extension of orfamide A is crucial for eliciting a specific Ca^2+^ signal in *C. reinhardtii*, or they could indicate that changes in compound structure may address different Ca^2+^ signalling pathways of which only a subset causes deflagellation. This hypothesis is in agreement with the observed bimodal Ca^2+^ release, in which the first peak is responsible for the deflagellation. Moreover, as mentioned before, stress factors such as high salt are known to modulate Ca^2+^ signatures, but do not cause deflagellation.^10^ The availability of more than 40 predicted Ca^2+^ channels in the membranes of *C. reinhardtii* also supports the above-mentioned conjecture.^16^

Therefore, we have analysed Ca^2+^ channels in *C. reinhardtii* in more detail. Most of them belong to the transient receptor potential (TRP)-type.^16^ Notably, TRP channels seem to be absent in land plants^16,17^ while they are found in mammals, yeast, algae and unicellular organisms.^18^ Some TRP channels in *C. reinhardtii* share key features with a sensory transduction-associated TRP channel in mammals^17^ and influence algal motility.^19^ TRP channels were originally grouped into seven subfamilies: TRPC (canonical), TRPV (vanilloid), TRPM (melastatin), TRPA (ankyrin), TRPN (NOMPC), TRPP (polycystic) and TRPML (mucolipin).^20^ Moreover, further subfamilies were recently classified, including TRPY/F (fungus-specific TRP channels), TRPS (soromelastanin) and TRPVL (vanilloid like).^21^ In order to classify the *C. reinhardtii* TRP channels, relate them to the known TRP families and get an overview of the so far known *C. reinhardtii* TRP channels, we conducted a comprehensive phylogenetic analysis (Fig. 3a; see details in Supporting Information). Interestingly, several of the green algal channels created clades of their own. TRP3, 5, 6, 7, 8, 10, 11, 16, 21, 22, 24 and 27 represented a clade related to the TRPV family. Another clade formed as a sister group to the TRPY/F channels (TRP12, 13, 15, 23, 26, 29 and 30). The data did not allow assigning TRP1 and TRP28 to a specific clade (Fig. 3a), but it seems that TRP28 shares a common ancestor with the families PKDREJ/TRPA/TRPV/TRPVL and TRP1 shares a common ancestor with the TRPC/TRPN/TRPS/TRPM families. TRP1 has been characterized and it has proven to show characteristics of different families as well as unique traits like a lipid binding pocket.^22^

**Fig. 3.**
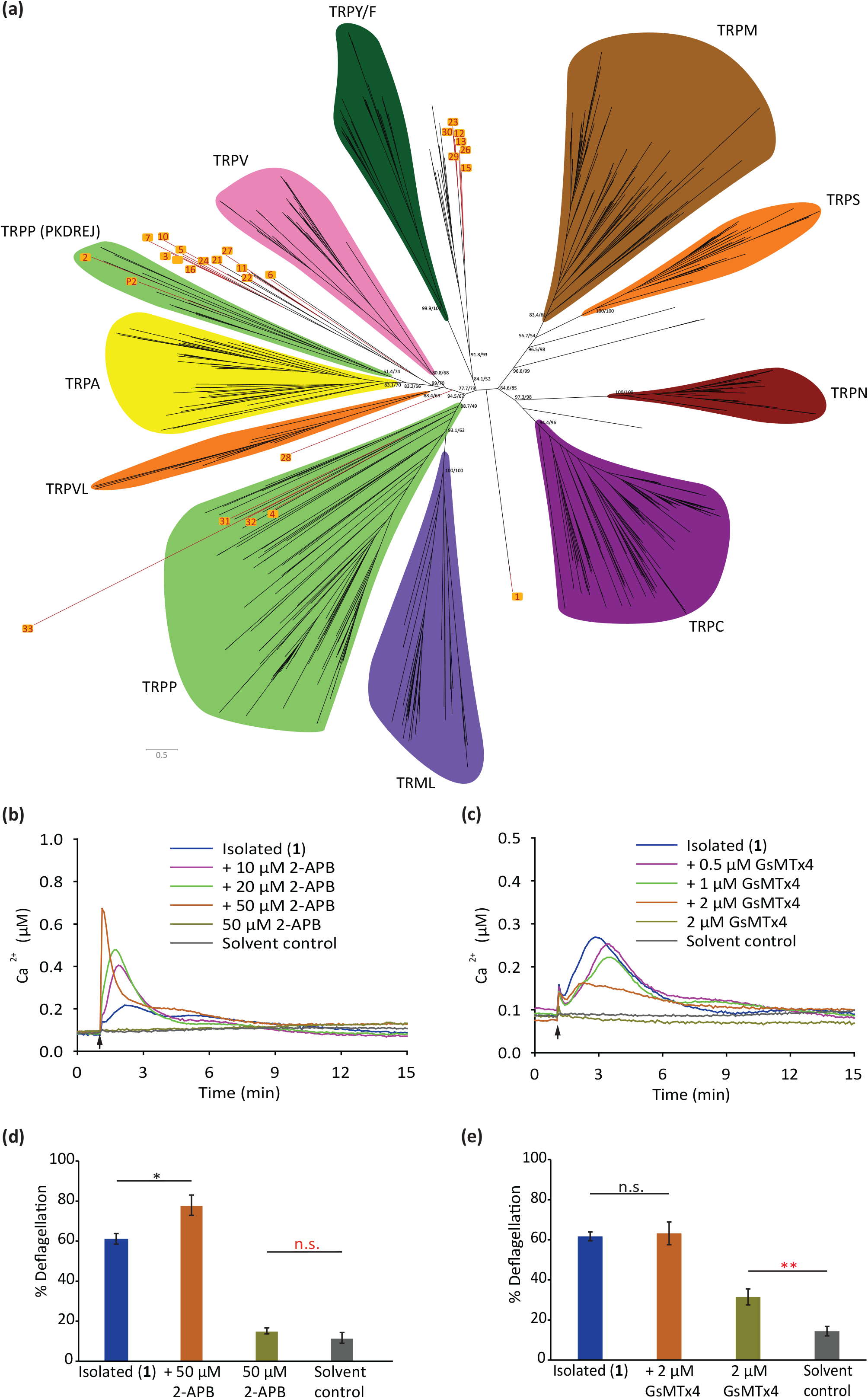
Phylogeny of TRP channels in *C. reinhardtii* ( a) and t he use of TRP channel modulators (b-e). (a) Phylogeny of TRP channels in *C. reinhardtii* (Table S2 in ESI). Phylogenetic tree containing the transmembrane domain of about 650 sequences, including 35 TRP channels with known structure (see ESI, Alignment Fasta and Table S3).^18^ The different families are highlighted. *C. reinhard ii* TRP channels are high-lighted in red. (b, c) Ca^2+^ signalling was studied with 5 μM of isolated orfamide A in the absence or presence of TRP channel modulators. Algal cells were treated with indicated concentrations of 2-APB and GsMTx4 for 1 min before the background measurements. Each line represents the mean of three independent biological replicates, and each biological replicate includes three technical replicates. (d, e) Proportions of deflagellated cells using 5 μM of isolated orfamide A in the absence or presence of TRP channel modulators. Algal cells were treated with the indicated concentration of 2-APB and GsMTx4 for 6 min before 5 μM of orfamide A were added. See legend of Figure 1 for details.

Notably, in our phylogenetic analysis, the TRPP family splits into two groups, one containing all of the polycystic kidney disease (PKD1-like and PKD2-like) and Brivido-related channels (which we will refer to as TRPP family), and a separate group containing the PKDREJ (polycystic kidney disease and receptor for egg jelly related protein)-like TRPP,^23^ which re-clustered as a sister group of the TRPA family. *C. reinhardtii* seems to have four channels belonging to the TRPP family (TRP4, 31, 32 and 33) and two channels of the PKDREJ family (TRP2 and TRPP2) (Fig. 3a).

To get a first insight into possibly involved Ca^2+^ channels, we applied two Ca^2+^ channel modulators of different selectivity as tool compounds and investigated their effect on orfamide-A-mediated Ca^2+^ signatures in *C. reinhardtii* together. At first, 2-aminoethoxydiphenyl borate (2-APB) was studied, a compound that blocks a subset of TRP channels while activating the thermosensitive channels TRPV1, TRPV2, TRPV3.^24^ 2-APB has been used before to characterize TRPV-type channels related to flagellar motility in *C. reinhardtii*.^19^ In the presence of this modulator, the Ca^2+^ signature of orfamide A became strongly elevated especially with regard to the first Ca^2+^ peak. The deflagellation rate was significantly increased (Fig. 3b, d), suggesting the involvement of TRPV-related channels in the orfamide-A-mediated signalling response. Alternatively, the peptide inhibitor GsMTx-4 was used. It inhibits mechanosensory channels (MSCs) and affects TRPC1.^25^ It has been applied in *C. reinhardtii* for the inhibition of mechanosensitive ion channels before.^10^ This tool compound strongly reduced the Ca^2+^ signature of orfamide A with regard to the second orfamide-A-induced Ca^2+^ peak, while the first one remained unaffected (Fig. 3c). In this case, the deflagellation rate was not significantly changed (Fig. 3e) corroborating the primary influence of the first Ca^2+^ peak on deflagellation (see also video 1). This observation conforms with the fact that these channels are also required for hypo-osmotic stress response, and that they do not cause deflagellation.^10^ Taken together, the tool compound and inhibitor studies suggest the involvement of multiple Ca^2+^ channels in orfamide-A-triggered signalling, involving TRPV- and TRPC-like channels, but also MSCs.

## Conclusions

Using synthetic variations of orfamide A and stereochemically probing its conformation, we have shown that the linear part of this CLiP is highly relevant for the specificity of the Ca^2+^ signalling event that causes deflagellation. The three *N*-terminal amino acids of the linear part as well as the terminal fatty acid region play important roles. Intriguingly, molecular editing of orfamide A indicated the triggering of at least two distinct Ca^2+^-signalling pathways, one that is apparently necessary for strong deflagellation (Glu2 and Thr3 changes), while the other still causes an increase in cytosolic Ca^2+^ in the algal cells when triggered but does not cause substantial deflagellation (Leu1 change). This effect is also corroborated by the relevance of the fatty acid hydroxylation. First data obtained with tool compounds suggest the involvement of different TRP-type and MSC Ca^2+^ channels in these reaction cascades, by showing synergy or antagonism in the Ca^2+^ response. Research toward a more detailed characterization of the Ca^2+^-channels in *C. reinhardtii* and toward elucidating the molecular mechanism responsible for acute deflagellation is currently ongoing.

## Supporting information

Aligment

## Author Contributions

M.M. and H.-D.A. conceived and designed the project; Y.B. and B.S. performed the chemistry experiments, Y.H., D.C.F. and V.H. performed the chemical biology experiments; D.C.F. performed the phylogenetic analysis; Y.H., Y.B., D.C.F. V.H. and B.S. analyzed the chemistry and chemical biology experiments; M.M. and H.-D.A. analyzed the data, with input of all authors; M.M. and H.-D.A. co-wrote the paper, with input from all authors.

## Conflicts of interest

There are no conflicts to declare.

## Acknowledgements

Y.B. was recipient of a predoctoral fellowship grant by the YOSHIDA foundation (JPN), V.H. received a fellowship of the International Leibniz Research School ILRS (under the head of the Jena School for Microbial Communication, JSMC) and D.C.F. had received a JSMC fellowship. M.M. and H.-D.A. received funding from the Deutsche Forschungsgemeinschaft (German Research Foundation) within the collaborative research center SFB1127/2 ChemBioSys—Project ID 239748522.

## Electronical Supplemental Information (ESI)

### Videos on cell motility upon treatment with orfamide A and variants

**Video 1** Synthetic orfamide A (**1**) - Ca^2+^ signals and cell motility are synchronized and shown

**Video 2** Commercial orfamide A (**1**) - Ca^2+^ signals and cell motility are synchronized and shown

**Video 3** Enantiomer of orfamide A (***ent*-1**) - Ca^2+^ signals and cell motility were synchronized and shown

**Video 4** Orfamide A variant (**5**) – Only cell motility is shown

**Video 5** Orfamide A variant (**7**) - Ca^2+^ signals and cell motility were synchronized and shown

**Video 6** Orfamide A variant (**2**) - Ca^2+^ signals and cell motility were synchronized and shown

## Supporting information

1. **Additional Figures**
2. **Chemical Methods including Table S1**
3. **Copies of NMR spectra**
4. **Biological Methods including Table S2**
5. **References SI**

Figure S1 Scheme of the dipeptide coupling followed by cyclization and side chain elongation Figure S2 Scheme of resin esterification followed by cyclization and terminal acylation

Figure S3 HPLC traces of orfamide A (1), ent-orfamide A (ent-1) and a mixture of 1 and ent-1 (220 nm, Condition: RP-A).

Figure S4 HPLC traces of orfamide A (1), *ent*-orfamide A (*ent*-1) and a mixture of 1 and *ent*-1 (205 nm, Condition: RP-A).

Figure S5 HPLC traces on chiral phase of orfamide A (1), *ent*-orfamide A (*ent*-1) and a mixture of 1 and

*ent*-1. (220 nm, Condition: RP-C, * artifacts).

Figure S6 HPLC traces on chiral phase of orfamide A (1), *ent*-orfamide A (*ent*-1) and a mixture of 1 and

*ent*-1. (205 nm, Condition: RP-C)

Table S1 One cycle of the SPPS using automated peptide synthesizer (Syro ll)

Table S2 Predicted TRP channels present in *Chlamydomonas reinhardtii* according to literature.

**Excel Table S3** Accession numbers and references where the protein was used for all sequences from the phylogenetic tree

**Alignment Fasta** Multiple sequence alignment of the transmembrane domain used to build the phylogenetic tree

## Notes

### Competing Interest Statement

The authors have declared no competing interest.

