## Supporting Information for "Orfamide-A-mediated bacterial-algal interactions involve specific Ca^2+^ signalling pathways"

### 1. Additional figures

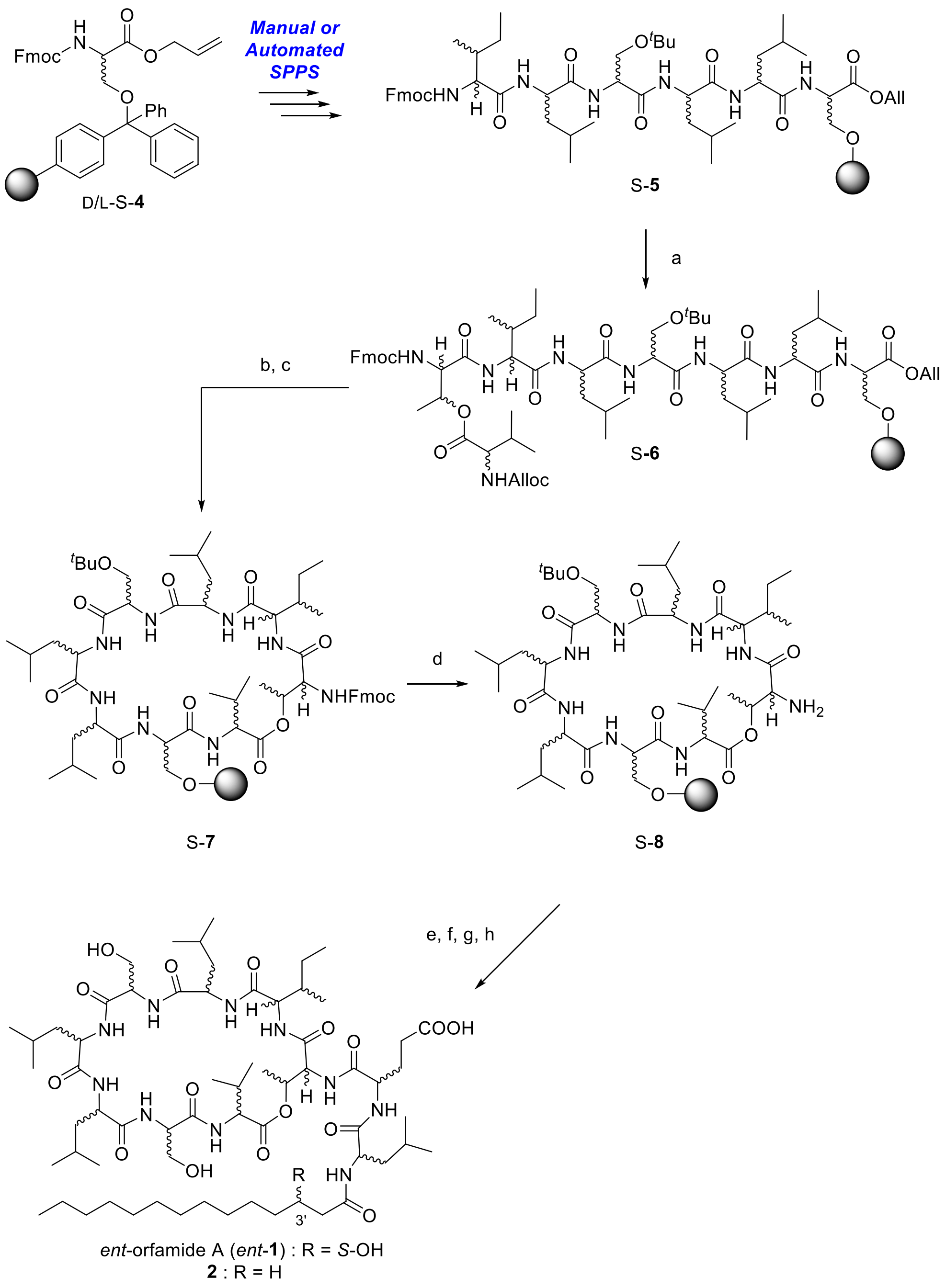


Reagents and Conditions: a) Fmoc-D-*allo*-Thr(Alloc-L-Val)-OH (*ent*-S-**3**) or Fmoc-L-*allo*-Thr(Alloc-D-Val)-OH (S-**3**), HATU, HOAt, collidine, DMF, rt; b) *cat.* $Pd(PPh_3)_4$, $PhSiH_3$, $CH_2Cl_2$, rt; c) HATU, HOAt, collidine, DMF, rt; d) 2% DBU 2% piperidine/DMF, 30 sec. x 2; e) Fmoc-D/L-Glu(O$^t$Bu)-OH, HBTU, HOBt, DIEA, DMF, rt; 20% piperidine/DMF; f) Fmoc-D/L-Leu-OH, HBTU, HOBt, DIEA, DMF, rt; 20% piperidine/DMF; g) (*S*)-3-TBSoxy tetradecanoic acid or tetradecanoic acid, HBTU, HOBt, DIEA, DMF, rt; h) 0.1 N HCl/HFIP, + 1% TIS, rt.

**Figure S1.** Scheme of the dipeptide coupling followed by cyclization and side chain elongation.

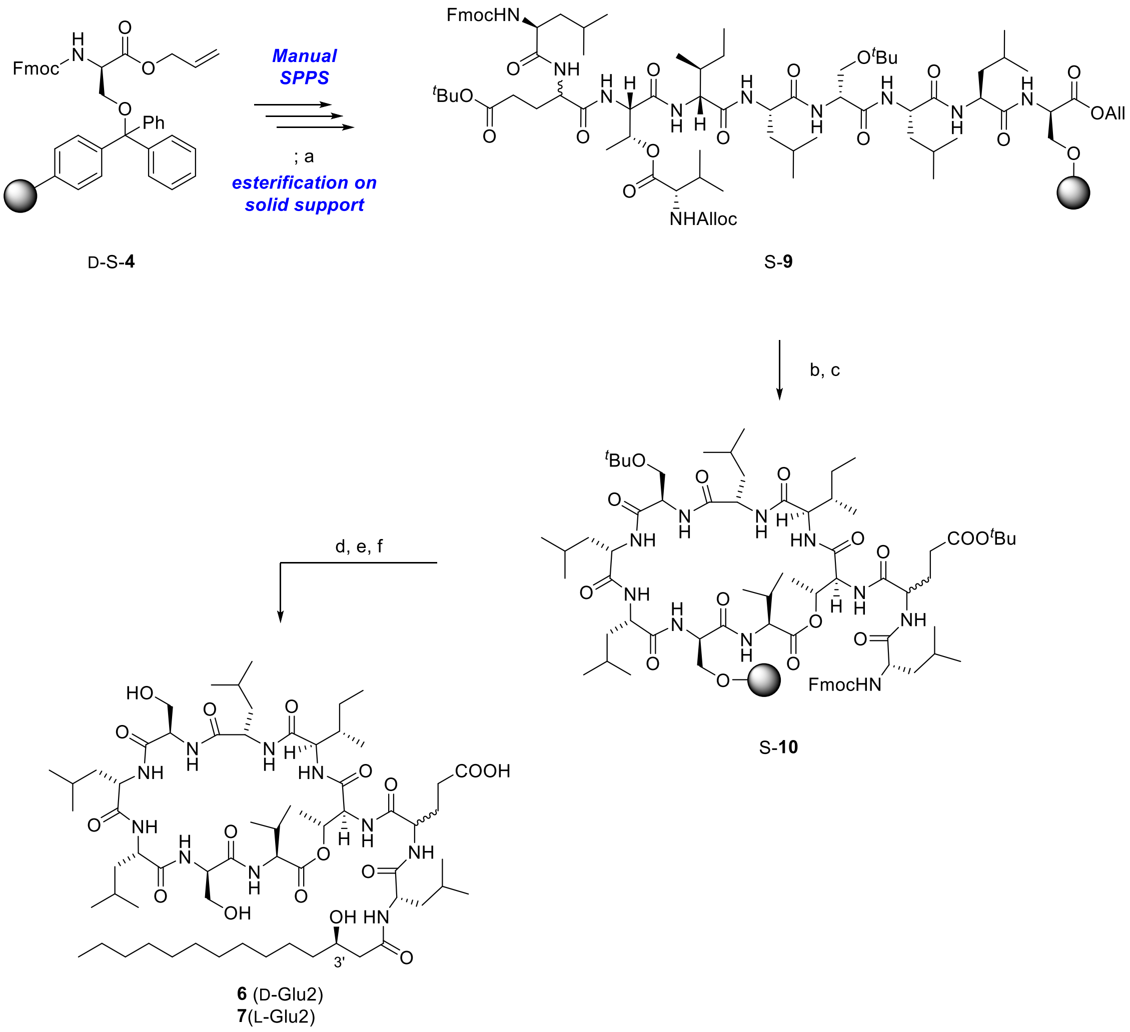


Reagents and conditions: a) Alloc-L-Valine, DIC, DMAP, THF, 40 °C or Alloc-L-Valine, DIC, DMAP, DMF, rt; b) *cat.* $Pd(PPh_3)_4$, $PhSiH_3$, $CH_2Cl_2$, rt; c) HATU, HOAt, DIEA, DMF, rt (Cyclization condition B); d) 20% piperidine/DMF, rt; e) (*R*)-3-hydroxytetradecanoic acid, HBTU, HOBt, DIEA, DMF, rt; f) 0.1 N HCl/HFIP, rt.

**Figure S2.** Scheme of resin esterification followed by cyclization and terminal acylation.

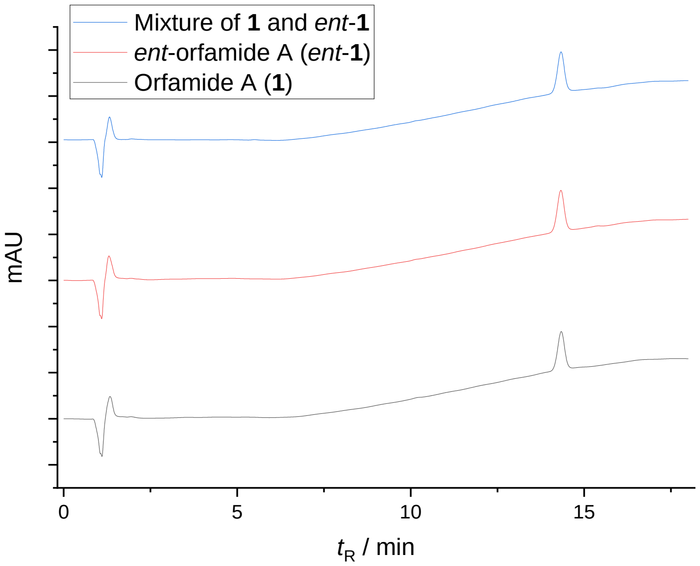


**Figure S3.** HPLC traces of synthetic orfamide A (**1**), *ent*-orfamide A (***ent*-1**) and a mixture of **1** and ***ent*-1** (220 nm, Condition: RP-A).

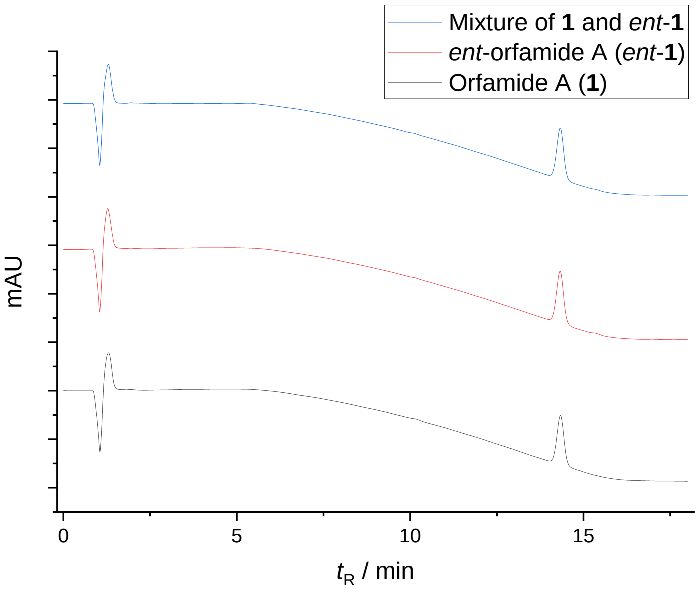


**Figure S4.** HPLC traces of synthetic orfamide A (**1**), *ent*-orfamide A (***ent*-1**) and a mixture of **1** and ***ent*-1** (205 nm, Condition: RP-A).

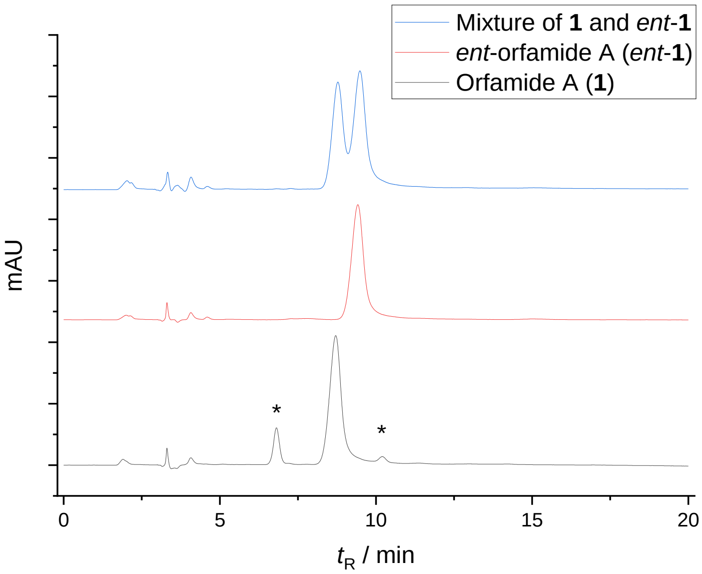


**Figure S5.** HPLC traces on chiral phase of synthetic orfamide A (**1**), *ent*-orfamide A (***ent*-1**) and a mixture of **1** and ***ent*-1**. (220 nm, Condition: RP-C, * artifacts).

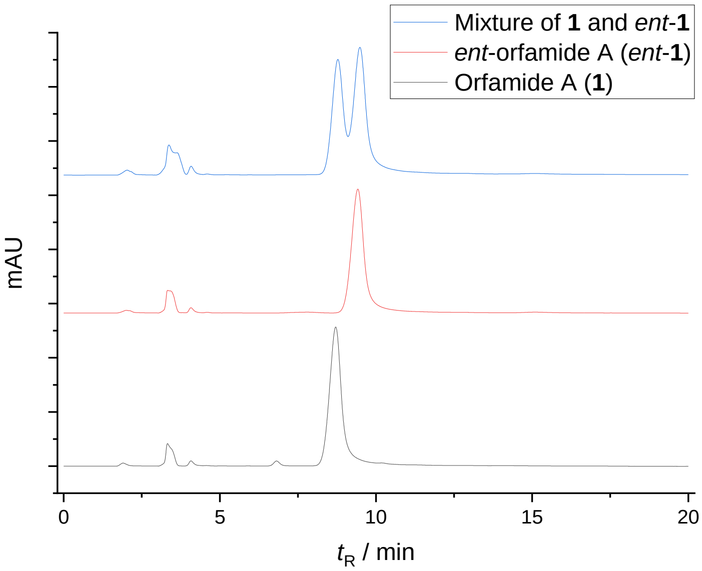


**Figure S6.** HPLC traces on chiral phase of synthetic orfamide A (**1**), *ent*-orfamide A (***ent*-1**) and a mixture of **1** and ***ent*-1**. (205 nm, Condition: RP-C).

### 2. Chemical Methods

#### 2.1 List of chemical abbreviations

| | |
|---|---|
| A.A. | Amino acid |
| Alloc | Allyloxycarbonyl |
| Bz | Benzoyl |
| DIC | *N,N′*-Diisopropylcarbodiimide |
| DIEA | *N,N*-Diisopropylethylamine |
| DMAP | 4-Dimethylaminopyridine |
| DMF | *N,N*-Dimethylformamide |
| DMSO | Dimethylsulfoxide |
| Dpm | Diphenylmethyl |
| EDCI | *N*-Ethyl-*N′*-(3-dimethylaminopropyl)-carbodiimide hydrochloride |
| eq. | Stoichiometric equivalent |
| F.A. | Fatty acid |
| Fmoc | 9-fluorenylmethyloxycarbonyl |
| HATU | 1-[*bis*(dimethylamino)methylene]-1*H*-1,2,3-triazolo[4,5*b*]pyridinium3-oxid hexafluorophosphate |
| HBTU | *N,N,N′,N′*-Tetramethyl-*O*-(1*H*-benzotriazol-1-yl)uronium hexafluorophosphate |
| HFIP | Hexafluoro-2-propanol |
| HOAt | 1-Hydroxy-7-azabenzotriazole |
| HOBt | 1-Hydroxybenzotriazole |
| SPPS | Solid phase peptide synthesis |
| TFA | Trifluoroacetic acid |
| TIS | Triisopropylsilane |
| THF | Tetrahydrofuran |

#### 2.2 General Synthesis Methods

##### 2.2.1 Reagents and reaction conditions

All reagents were purchased from Acros Chemicals, Alfa Aesar, ABCR, Carbolution Chemicals, Carbosynth, Fischer Chemical, fluoroChem, GL Biochem (Shanghai), GRÜSSING, Manchester Organics, Merck, Novabiochem, Sigma-Aldrich, TCI Europe and VWR. All solvents, if not purchased in purity or dryness suitable, were distilled. THF was heated to reflux with Na(0) pieces and benzophenone under $N_2$ atmosphere until a blue-purple color persisted, and distilled. $CH_2Cl_2$ was heated to reflux under $N_2$ atmosphere for 1h with $CaH_2$, and distilled. DMF and MeOH (HPLC grade) were stored with 3Å molecular sieves at least for 24 h prior to use. All solvents for flash chromatography were distilled by using a rotary evaporator prior to use. If necessary, solvents were degassed by purging with $N_2$ or Ar for at least 15 minutes. Deionized water was used for all experiments. All reactions were performed under protective atmosphere ($N_2$ or Ar) if not stated otherwise.

#### 2.2.2 Thin layer chromatography (TLC)

Merck precoated silica gel plates (60 $F_{254}$) were used. Compounds were visualized by using ultraviolet light irradiation at 254 nm and/or by using the following staining agent (dip, dry & heat development).

Phosphomolybdic acid (PMA): $12MoO_3 \cdot H_3PO_4$ (5 g) in EtOH (100 mL).

#### 2.2.3 Silica gel flash liquid chromatography

Compound purification was performed using silica gel from MACHEREY-NAGEL (particle size 40-60 µm) under approximately 0.2-0.4 bar pressure, applying apdapted conditions of Still et. al.[1]

#### 2.2.4 NMR spectroscopy

$^{1}$H- and $^{13}$C-NMR spectra were recorded using Bruker Advance I 250, Fourier 300, Advance III 400, Advance III HD 500 or Advance III 600 system. Spectra were calibrated to appropriate residual solvent peaks (chloroform-d, methanol-$d_4$, DMSO-$d_6$).[2]

#### 2.2.5 Analytical reverse phase HPLC (RP-HPLC)

Analyses were performed on a SHIMADZU system consisting of a system controller (SLC- 10A VP), a column oven (CTO-10AC VP), an auto-injector (SIL-10ADVP), a degasser (DGU- 14A), three pumps (LC-10AT VP), a diode array detector (SPD-M20A), a fluorescence detector (RF-10AXL) and an analytical column. The column was equilibrated to starting condition of each method prior to sample injections.

Eluent System: A = acetonitrile, B = water, C = 2% TFA in water, Flow rate: 1 mL/min, Column oven: 25 °C, Detection: diode array 190-800 nm.

**Condition A (RP-A)**

Column: MACHEREY-NAGEL NUCLEODUR C18 Gravity, 5 µm, 125 × 4 mm.

Gradient: Eluent A: 70% (1 min), 70-95% (10 min), 95% (5 min), 95-70% (0.2 min), 70% (5.8 min).
Eluent C: 5% (22 min).

Flow rate: 1 mL/min.

**Condition B (RP-B)**

Column: MACHEREY-NAGEL NUCLEODUR C18 Gravity, 5 µm, 125 × 4 mm.

Gradient: Eluent A: 70% (1 min), 70-95% (10 min), 95% (8 min), 95-70% (0.2 min), 70% (5.8 min).
Eluent C: 5% (25 min).

Flow rate: 1 mL/min

**Condition C (RP-C)**

Column: DAICEL CHIRALPAK® AD-RH, 5 µm, 150 × 4.6 mm (Chiral column).

Gradient: Eluent A: 45% (20 min), Eluent C: 0% (20 min).

Flow rate: 0.6 mL/min.

#### 2.2.6 Reverse phase preparative HPLC (RP-Prep. HPLC)

Purifications of final products were performed on a SHIMADZU system consisting of a system controller (SLC- 10A VP), two pumps (LC-8A), a UV-VIS detector (SPD-10AVP) and a fraction collector (FRC-10A). Columns were equilibrated to starting conditions of each method prior to sample injections.

Eluent System: A = 0.1% TFA in acetonitrile, B= 0.1% TFA in water, Detection: UV-VIS 220 nm.

**Condition A (Prep.-A)**

Column: MACHEREY-NAGEL NUCLEODUR C18 Gravity, 5 µm, 250 × 16 mm.

Gradient: Eluent A: 60-95% (35 min), 95% (15 min), 95-60% (5 min), 60% (10 min).

Flow rate: 10 mL/min.

**Condition B (Prep.-B)**

Column: MACHEREY-NAGEL NUCLEODUR C18 Gravity, 5 µm, 250 × 16 mm.

Gradient: Eluent A: 70-80% (20 min), 80-95% (15 min), 95% (10 min), 95-70% (2 min), 70% (14 min).

Flow rate: 10 mL/min.

**Condition C (Prep.-C)**

Column: MACHEREY-NAGEL NUCLEODUR C18 Gravity, 5 µm, 250 × 16 mm.

Gradient: Eluent A: 90-95% (10 min), 95% (27 min), 95-90% (10 min), 90% (13 min).

Flow rate: 10 mL/min.

**Condition D (Prep.-D)**

Column: KNAUER Eurospher 100-5 C8, 250 × 16 mm.

Gradient : Eluent A: 80% (3 min), 80-85% (10 min), 85% (5 min), 85-95% (20 min), 95% (15 min), 95-80% (2 min), 80% (10 min).

Flow rate: 20 mL/min.

#### 2.2.7 Liquid chromatography mass spectrometry (LC/MS)

Analyses were performed on a SHIMADZU system consisting of a system controller (SLC- 10A VP), a column oven (CTO-10AC VP), an auto-injector (SIL-10AD VP), a degasser (DGU- 14A), two pumps (LC-10AT VP), an analytical column (MACHEREY-NAGEL NUCLEODUR C18 Isis, 3 µm), a post-column flow splitter (Thermo scientific, ICP-04-20), a UV-VIS detector (SPD-10A VP) and a MS detector (Finnigan LCQ spectrometer). The column was equilibrated to starting conditions of each method prior to the sample injections.

Eluent System: A = 0.1% HCOOH in acetonitrile, B= 0.1% HCOOH in water, Flow rate: 1 mL/min, Detection: UV/VIS 220 or 254 nm, Column oven: 25 °C.

#### 2.2.8 Liquid chromatography high resolution mass spectrometry (LC/HRMS)

Analyses were performed on a Thermo scientific UltiMate 3000 UHPLC system consisting of a pump, an auto sampler, a column compartment, a diode array detector and a high resolution Q-TOF MS spectrometer (maXis Impact, BRUKER DALTRONICS, Bremen Germany). The column was equilibrated to starting conditions of each method prior to the sample injections.

Eluent System: A = 0.1% HCOOH in acetonitrile, B= 0.1% HCOOH in water, Flow rate: 0.5 mL/min, Detection: Diode array 200 - 400 nm.

#### 2.2.9 Specific optical rotation

Optical rotations were recorded in a Jasco P-2000 polarimeter at 589 nm. Path length of cuvettes was d = 10 mm. Concentrations (c) are given in g/100 mL.

#### 2.2.10 Melting points

Melting points were measured with Büchi B-545 melting point apparatus. One-side open capillaries were used.

### 2.3 Synthesis procedures and physical data

#### 2.3.1 Building blocks

(*R*) or (*S*)-3-((*tert*-butyldimethylsilyl)oxy)tetradecanoic acids were synthesized as reported.[3,4] Fmoc-L-*allo*-Thr(Alloc-D-Val)-OH (**S-3**) was prepared in a same procedure as its enantiomer.[3]

**Fmoc-L-*allo*-Thr-ODpm (S-1)**

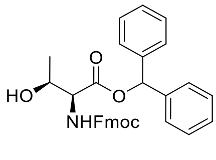


S-**1**

Fmoc-L-*allo*-Thr-ODpm (S-**1**) was prepared[3] from Fmoc-L-*allo*-Threonine (331 mg, 0.970 mmol) and was obtained as colorless solid (436 mg, 0.860 mmol, 89%).

**TLC:** $R_f$ = 0.42 (petroleum ether/EtOAc = 2:1).

**$^1$H-NMR (CDCl$_3$, 300 MHz, 297 K)** $\delta$ = 1.09 (d, $J$ = 6.4 Hz, 3 H), 2.81 (br s, 1 H), 4.06 - 4.30 (m, 2 H), 4.42 (m, 2 H), 4.60 (br dd, $J$ = 7.1, 3.1 Hz, 1 H), 5.72 (br d, $J$ = 7.4 Hz, 1 H), 6.94 (s, 1 H), 7.26 - 7.43 (m, 14 H), 7.59 (br d, $J$ = 7.3 Hz, 2 H), 7.77 (d, $J$ = 7.5 Hz, 2 H) ppm.

**$^{13}$C{$^1$H}-NMR (CDCl$_3$, 75 MHz, 297 K)** $\delta$ = 18.7, 47.2, 59.6, 67.5, 69.2, 77.4, 78.8, 120.1, 125.2, 125.2, 127.1, 127.2, 127.4, 127.8, 127.9, 128.3, 128.5, 128.8, 139.3, 139.3, 141.4, 141.4, 143.7, 143.8, 156.9, 169.5 ppm.

**HRMS (ESI-TOF)** calculated for $C_{32}H_{29}NO_5$ $[M+H]^+$ 508.2118; found 508.2124.

**$[\alpha]_D$ =** -11.7 ($CH_2Cl_2$, c = 1.0, 26.2 °C).

**Melting Point:** $T_m$ = 168.4 °C.

**Fmoc-L-allo-Thr(Alloc-D-Val)-ODpm (S-2)**

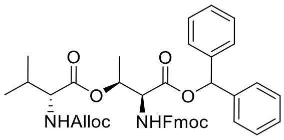


S-**2**

Fmoc-L-*allo*-Thr(D-Alloc-Val)-ODpm (**S-2**) was prepared from Fmoc-L-*allo*-Thr-ODpm (**S-1**) (200 mg, 0.394 mmol, 1.0 eq.) and Alloc-D-Valine (96.3 mg, 0.479 mmol, 1.2 eq.),[3] and was obtained as colorless solid (259 mg, 0.375 mmol, 95%).

**TLC:** $R_f$ = 0.73 (petroleum ether/EtOAc = 2:1).

**$^1$H-NMR (CDCl$_3$, 300 MHz, 297 K)** $\delta$ = 0.84 (d, $J$ = 6.8 Hz, 3 H), 0.90 (d, $J$ = 6.8 Hz, 3 H), 1.25 (d, $J$ = 6.7 Hz, 3 H), 2.03 (m, 1 H), 4.12 (m, 1 H), 4.22 (t, $J$ = 7.3 Hz, 1 H), 4.36 (br d, $J$ = 7.3 Hz, 2 H), 4.55 (dd, $J$ = 13.5, 6.1 Hz, 2 H), 4.77 (dd, $J$ = 8.6, 3.1 Hz, 1 H), 5.13 (m, 1 H), 5.23 (m, 1 H), 5.29 (m, 1 H), 5.35 (m, 1 H), 5.85 (m, 2 H), 6.99 (s, 1 H), 7.28 - 7.43 (m, 14 H), 7.61 (t, $J$ = 6.4 Hz (ent:8.2 Hz), 2 H), 7.76 (d, $J$ = 7.5 Hz, 2 H) ppm.

**$^{13}$C{$^1$H}-NMR (CDCl$_3$, 75 MHz, 297 K)** $\delta$ = 16.0, 17.6, 19.1, 31.0, 47.2, 57.4, 59.3, 66.0, 67.7, 72.1, 78.9, 118.0, 120.1, 125.3, 127.1, 127.2, 127.6, 127.9, 128.4, 128.5, 128.8, 128.8, 132.7, 139.2, 139.3, 141.4, 143.9, 156.2, 156.4, 168.1, 171.5 ppm.

**HRMS (ESI-TOF)** calculated for $C_{41}H_{42}N_2O_8$ $[M+H]^+$ 691.3014; found 691.3034.

**$[\alpha]_D$** = -8.0 ($CHCl_3$, c = 1.0, 25.9 °C).

**Melting Point:** $T_m$ = 40 - 60 °C.

**Fmoc-L-allo-Thr(Alloc-D-Val)-OH (S-3)**

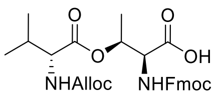


S-**3**

Fmoc-L-*allo*-Thr(D-Alloc-Val)-OH (S-**3**) was prepared[3] from Fmoc-L-*allo*-Thr(D-Alloc-Val)-ODpm (**S-2**) (600 mg, 0.869 mmol) and was obtained as colorless solid (452 mg, 0.862 mmol, 99%).

**TLC:** $R_f$ = 0.54 ($CH_2Cl_2$/MeOH = 20:1 + 1% HCOOH).

**$^1$H-NMR ($CDCl_3$, 300 MHz, 297 K)** $\delta$ = 0.89 (d, $J$ = 7.0 Hz, 3 H), 0.95 (d, $J$ = 7.0 Hz, 3 H), 1.42 (m, 3 H enantiomer:1.48, d), 2.13 (m, 1 H), 4.18 (m, 1 H), 4.25 (m, 1 H), 4.37 (m, 2 H), 4.57 (m, 3 H), 5.15 (d, $J$ = 10.2 Hz, 1 H), 5.24 (d, $J$ = 17.2 Hz, 1 H), 5.37 (m, 1 H), 5.51 (m, 1 H), 5.81 (m, 1 H), 6.44 (br s, 1 H), 7.26 (m, 2 H), 7.36 (t, $J$ = 7.3 Hz, 2 H), 7.55 (t, $J$ = 7.9 Hz, 2 H), 7.73 (d, $J$ = 7.5 Hz, 2 H) ppm.

**$^{13}$C{$^1$H}-NMR ($CDCl_3$, 126 MHz, 297 K)** $\delta$ = 16.6, 17.4, 19.1, 31.0, 47.2, 57.7, 59.1, 66.5, 67.6, 72.6, 118.3, 120.0, 125.4, 127.2, 127.8, 132.3, 141.3, 143.9, 144.0, 156.4, 157.1, 171.5 ppm.

**HRMS (ESI-TOF)** calculated for $C_{41}H_{42}N_2O_8$ $[M+H]^+$ 525.2231; found 525.2238.

$[\boldsymbol{\alpha}]_{\boldsymbol{D}}$ = 20.6 ($CHCl_3$, c = 1.0, 25.5 °C).

**Melting Point:** $T_m$ = 85°C.

#### 2.3.2 General procedures for solid phase peptide synthesis[3]

All peptide syntheses were performed on solid support. Reactions were monitored by small scale cleavage of the peptidyl resin using the following conditions and subjecting the supernatant to LC/MS and/or RP-HPLC analyses. Amounts of reagents and solvents used were calculated based on the initial amino acid loading on resin. Yields were calculated based on resin loading.

Condition A: 5% TFA in $CH_2Cl_2/H_2O$ (2.5:1; v/v), rt, 15 min.

Condition B: 0.1 N HCl/HFIP (10 µL 37% aq. HCl / 990 µL HFIP) + 1% TIS, rt, 30 – 60 min.[4,5]

#### 2.3.3 Fmoc-D/L-Ser(OH)-OAll (S-4) loading on trityl chloride resin

The mixture of Fmoc-D/L-Ser(OH)-OAll (S-4) (0.5 - 1.5 eq.), trityl chloride resin (1.64 mmol/g; 1 eq.) and pyridine (3 eq.) in dry THF (5 - 10 mL/g dry resin) were stirred at 70 °C for 7 h. The resin was washed with $CH_2Cl_2$ (6 ×) and then treated with a solution of $CH_2Cl_2$/MeOH/DIEA (17:2:1, v/v/v) for 20 min. After the removal of solvents the resin was washed with $CH_2Cl_2$ (6 ×) and MeOH (1 ×), and dried in vacuo overnight. The loading was determined by a small scale Fmoc cleavage using 20% piperidine/DMF (v/v) and measurement of the absorption of the cleaved piperidine-dibenzofulvene adduct.

#### 2.3.4 SPPS using an automated parallel peptide synthesizer

An automated peptide synthesizer (MultiSyn Tech, Syro II) was programmed to couple amino acids sequentially from C-terminus to N-terminus for **S-5** synthesis. A single coupling cycle consisted of Fmoc deprotection, amino acid coupling and washing with DMF (3 ×) after each step. All steps were performed at room temperature in open air. The detailed protocol used to program the peptide synthesizer is described in Table S1.

**Table S1: One cycle of the SPPS using automated peptide synthesizer (Syro ll)**

| Step | Operation | Solvent/Reagent | Time (repetitions) |
|---|---|---|---|
| 1 | Swelling | DMF | 20 min 30 s (2 ×) [a, b] |
| 2 | Fmoc deprotection | 20 % piperidine in DMF (v/v) | 3 min (1 ×) [a, c]<br>7 min (2 ×) [a, c] |
| 3 | Wash | DMF | 3 min (3 ×) [a, c] |
| 4 | Coupling | Fmoc-A.A. solution in DMF (0.4 M, 4 eq.); add HBTU/HOBt in DMF (0.3 M each, 4 eq.); add DIEA in NMP (1.3 M) | 2 h* (1 ×) [a, d] |
| 5 | Wash | DMF | 3 min (3 ×) [a, c] |

[a] Vortex = 20 sec.; [b] Break = 3 min; [c] Break = 1 min; [d] Break = 5 min.

* Couplings of Fmoc-L-*allo*-Isoleucine were performed for 8 h**.**

#### 2.3.5 Manual SPPS

The resin was swollen prior to all steps.

#### 2.3.6 Fmoc deprotection conditions

Fmoc deprotection was performed under two conditions.

**Condition A:**

The deprotection solution (20% piperidine/DMF (v/v)) was added to the peptidyl resin in a syringe and the syringe was shaking for 5 min at room temperature. The Fmoc deprotection solution was removed, and the deprotection step was repeated once for another 15 min, and the resulting peptidyl resin was washed with DMF (6 ×) prior to the subsequent coupling.

**Condition B:**

The deprotection solution (DBU/piperidine/DMF (2/2/96; v/v/v)) was added to the peptidyl resin in a syringe and the syringe was shaking for 30 seconds at room temperature. The Fmoc deprotection solution was removed and the resin was washed with DMF (1 ×). The deprotection step was repeated once for another 30 seconds, and the resulting peptidyl resin was washed with DMF (6 ×). All procedures were performed within 5 min prior to the next step.

#### 2.3.7 Amino acid/building block coupling conditions

All couplings were performed at room temperature.

**Coupling cocktail A:**

Carboxylic acid (A.A.) or building block (4 eq.), HBTU (4 eq.), HOBt (4 eq.), DIEA (8 eq.) in DMF (0.25 M; with respect to A.A. or building blocks).

**<u>Coupling cocktail B</u>:**

A.A. or building block (5 eq.), HBTU (5 eq.), HOBt (5 eq.), DIEA (10 eq.) in DMF (0.4 M; with respect to A.A. or building blocks).

**<u>Coupling cocktail C</u>:**

A.A. or building block (2 eq.), HBTU (2 eq.), HOBt (2 eq.), DIEA (4 eq.) in DMF (0.4 M; with respect to A.A. or building blocks).

**<u>Coupling cocktail D</u>:**

A.A. or building block (2 eq.), HATU (2 eq.), HOAt (2 eq.), collidine (4 eq.) in DMF (0.4 M; with respect to A.A. or building blocks).

#### 2.3.8 Manual SPPS for S-5 and S-9

The corresponding amino acids were coupled to the peptidyl resin under the conditions described below.

Fmoc deprotection: Condition A.

Fmoc-A.A.: Coupling cocktail A, 2 h at room temperature.

#### 2.3.9 Dipeptide coupling (Figure S1)

After Fmoc deprotection (Condition A), the coupling cocktail D with dipeptide (S-3 or *ent*-S-3) was added to the syringe containing the peptidyl resin, and the syringe was shaken for 1 h at room temperature. The coupling mixture was removed from the syringe and the peptidyl resin was washed with DMF (6 ×) and $CH_2Cl_2$ (3 ×).

#### 2.3.10 On-resin esterification (Figure S2)

The on-resin esterification was performed under one of the conditions described below.

**<u>Condition A (On resin esterification with L-Glu2 derivative)</u>:**

To the reaction flask containing peptidyl resin was added Alloc-L-Valine (10 eq.) and DMAP (1 eq.) in THF and DIC (10 eq.) and the mixture was stirred at 40 °C for 4 h. Then additional DIC (10 eq.) was added to the reaction mixture and stirred for another 4 h. The peptidyl resin was washed with $CH_2Cl_2$ (6 ×), DMF (3 ×) and $CH_2Cl_2$ (6 ×). This step was repeated if necessary.

**<u>Condition B (On resin esterification with D-Glu2 derivative)</u>:**

To the syringe containing peptidyl resin was added a solution of Alloc-L-Valine (10 eq.), DIC (10 eq.) and DMAP (1 eq.) in DMF and the syringe was shaken for 4 h at room temperature. The coupling mixture was removed and the peptidyl resin was washed with DMF (6 ×) and $CH_2Cl_2$ (3 ×). This step was repeated if necessary.

#### 2.3.11 Alloc/all deprotection

Peptidyl resin was dried under high vacuum prior to Alloc/All deprotection and the resin was swollen in $CH_2Cl_2$ under Ar atmosphere. A solution of $Pd(PPh_3)_4$ (0.2 eq.) and $PhSiH_3$ (20 eq.) in $CH_2Cl_2$ (4.05 mM with respect to $Pd(PPh_3)_4$) was added to the reaction pot containing the resin and the mixture was stirred at room temperature for 2 h in the dark. The resin was washed with $CH_2Cl_2$ (3 ×), DMF (3 ×) and $CH_2Cl_2$ (3 ×).

#### 2.3.12 On-resin cyclization

The on-resin cyclization was performed under one of the conditions described below.

**Condition A (Figure S1):**

To the syringe containing peptidyl resin was added a solution of HATU (2 eq.), HOAt (2 eq.) and collidine (4 eq.) in DMF (0.4 M with respect to HATU) and the syringe was shaken for 1 h. The reagents mixture was removed then the peptidyl resin was washed with DMF (6 ×) and $CH_2Cl_2$ (3 ×).

**Condition B (Figure S2):**

To the syringe containing peptidyl resin was added a solution of HATU (5 eq.), HOAt (5 eq.) and DIEA (10 eq.) in DMF (0.25 M with respect to HATU) and the syringe was shaken for 3 - 18 h. The reagent mixture was removed then the peptidyl resin was washed with DMF (6 ×) and $CH_2Cl_2$ (3 ×).

#### 2.3.13 Rapid Fmoc deprotection; Subsequent Amino acid coupling (Figure S1)

After cyclization, the Fmoc group of cyclic octadepsipeptide **S-7** was quickly removed (Fmoc cleavage condition B) and the resulting peptidyl resin was subjected to subsequent A.A. coupling (Coupling cocktail B). The mixture of Fmoc-D/L-Glu(O$^t$Bu)-OH, HBTU and HOBt in DMF was stirred for 20 min prior to the coupling, and DIEA was added to the mixture just before the coupling cocktail was added to the syringe. The mixture was shaken for 1 h at room temperature. The reagents mixture was removed then the peptidyl resin was washed with DMF (6 ×) and $CH_2Cl_2$ (3 ×).

#### 2.3.14 Side chain elongation

The corresponding amino acids and fatty acids were coupled to the peptidyl resin under the conditions described below.

| | |
|---|---|
| Fmoc deprotection: | Condition A. |
| Fmoc-A.A.: | Coupling cocktail A, 2 h, at room temperature. |
| Fatty acid: | Coupling cocktail C, 2 - 16 h, at room temperature. |

#### 2.3.15 Cleavage and global deprotection of peptide from resin

Peptidyl resin was dried under high vacuum prior to peptide cleavage from resin. Peptides were cleaved from resin by cleavage condition B (ca. 100 mg peptidyl resin/1 mL cleavage cocktail). The syringe was shaken for 20 min at room temperature and the solution was collected in a round bottom flask. This procedure was repeated three times. The resulting mixture was stirred further at room temperature until complete deprotection was observed by LC/MS. Then the mixture evaporated with toluene and the crude mixture was purified by RP-prep. HPLC.

### 2.4 Analytical data of synthesized cyclic lipodepsipeptides

For analytical data of orfamide A (1), compounds 3, 4, and 5, see Bando et al.[3]

#### 2.4.1 *ent*-Orfamide A (*ent*-1)

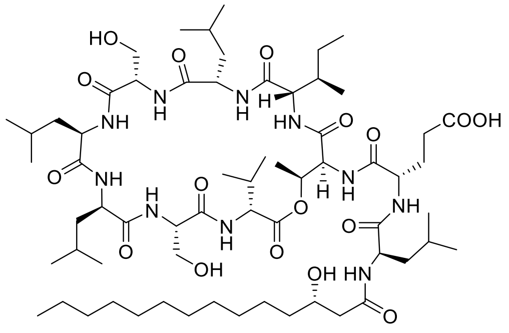


*ent*-orfamide A
(*ent*-**1**)

Obtained as colorless resin (11.6 mg, 8.95 µmol, 11% from resin loading 0.36 mmol/g). For HPLC-comparison with synthetic orfamide A, see figures S3-S6.

**RP-HPLC**: $t_R$ = 14.3 min (RP-A).

**RP-prep.HPLC**: Prep.-D.

**$^1$H-NMR (MeOH-$d_4$, 600 MHz, 297 K)** $\delta$ = 0.82 – 1.01 (m, 39 H), 1.20 (m, 1 H), 1.25 - 1.36 (m, 18 H), 1.38 (d, $J$ = 6.0 Hz, 3 H), 1.42 - 1.83 (m, 15 H), 2.01 (m, 2 H), 2.14 (m, 2 H), 2.36 (dd, $J$ = 14.2, 9.1 Hz, 1H), 2.45 (m, 3 H), 3.82 (dd, $J$ = 11.4, 4.5 Hz, 1 H), 3.87 (dd, $J$ = 11.6, 4.4 Hz, 1 H), 3.92 (m, 2 H), 3.98 (d, $J$ = 7.6 Hz, 1 H), 4.08 (m, 1 H), 4.13 (t, $J$ = 7.3 Hz, 1 H), 4.17 (dd, $J$ = 8.7, 6.1 Hz, 1 H), 4.22 (m, 1 H), 4.30 (m, 1 H), 4.34 (m, 4 H), 4.47 (m, 1 H), 5.37 (m, 1 H) ppm. (COOH, CONH and OH not assigned).

**HRMS (ESI-TOF)** calculated for $C_{64}H_{114}N_{10}O_{17}$ $[M+H]^+$ 1295.8436; found 1295.8433.

**2.4.2 3’-deoxyorfamide A (2)**

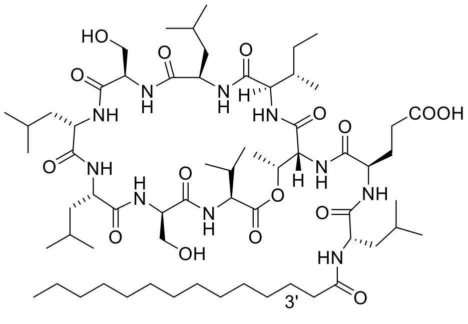

3'-deoxyorfamide A (**2**)

Obtained as colorless resin (7.5 mg, 5.86 µmol, 25% from resin loading).

**RP-HPLC:** $t_R$ = 17.9 min (RP-B).

**RP-prep.HPLC:** Prep.-C.

**$^1$H-NMR (MeOH-d$_4$, 600 MHz, 297 K)** *δ* = 0.83 (d, *J* = 6.8 Hz, 3 H), 0.85 - 1.02 (m, 36 H), 1.20 (m, 1 H), 1.25 - 1.39 (m, 20 H), 1.37 (d, *J* = 6.1 Hz, 3 H), 1.43 - 1.82 (m, 16 H), 2.01 (m, 2 H), 2.17 (m, 2 H), 2.30 (t, *J* = 7.6 Hz, 2 H), 2.45 (m, 2 H), 3.81 (dd, *J* = 11.4, 4.6 Hz, 1 H), 3.89 (m, 2 H), 3.94 (dd, *J* = 11.6, 5.7 Hz, 1 H), 3.99 (d, *J* = 7.5 Hz, 1 H), 4.12 (t, *J* = 7.4 Hz, 1 H), 4.18 (dd, *J* = 8.8, 5.9 Hz, 1 H), 4.23 (dd, *J* = 11.2, 3.9 Hz, 1 H), 4.27 - 4.39 (m, 5 H), 4.46 (m, 1 H), 5.39 (m, 1 H) ppm. (COOH, CONH and OH not assigned).

**HRMS (ESI-TOF)** calculated for $C_{64}H_{114}N_{10}O_{16}$ $[M+H]^+$ 1279.8487; found 1279.8491.

**2.4.3 Compound 6 (L-Thr3, L-Leu5 derivative)**

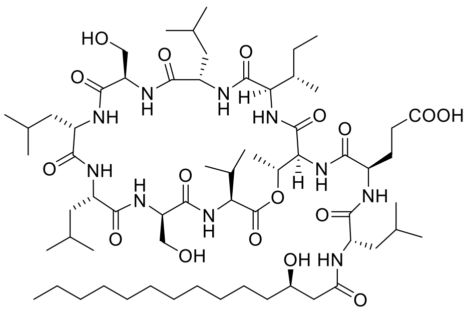

**6**

Obtained as colorless resin (8.1 mg, 6.25 µmol, 21% from a resin loading of 0.87 mmol/g).

**RP-HPLC:** $t_R$ = 6.9 min (RP-A).

**RP-prep.HPLC:** Prep.-B.

**$^1$H-NMR (MeOH-d$_4$, 600 MHz, 297 K)** *δ* = 0.85 - 1.06 (m, 39 H), 1.20 (m, 1 H), 1.23 (d, *J* = 9.7 Hz, 3 H), 1.25 - 1.39 (m, 18 H), 1.41 - 1.80 (m, 15 H), 1.83 (m, 1 H), 1.93 (m, 1 H), 2.26 (m, 2 H), 2.32 (m, 1 H), 2.36 (dd, *J* = 14.3, 8.2 Hz, 1 H), 2.45 (m, 2 H), 3.41 (m, 1 H), 3.48 (dd, *J* = 11.3, 5.7 Hz, 1 H), 3.85 (m, 2 H), 3.92 (dd, *J* = 11.4, 5.1 Hz, 1 H), 3.99 (m, 1 H), 4.19 (m, 1 H), 4.26 (t, *J* = 4.4 Hz, 1 H), 4.39 (m, 3 H), 4.53 (m, 2 H), 4.82 (m, 1 H), 4.98 (br s, 1 H), 5.46 (m, 1 H) ppm. (COOH, CONH and OH not assigned).

**HRMS (ESI-TOF)** calculated for $C_{64}H_{114}N_{10}O_{17}$ $[M+H]^+$ 1295.8436; found 1295.8441.

**2.4.4 Compound 7 (L-Glu2, L-Thr3, L-Leu5 derivative)**

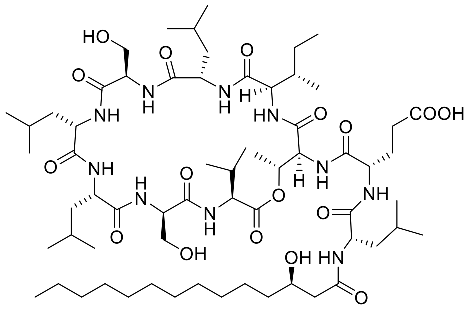

**7**

Obtained as colorless resin (4.2 mg, 3.24 µmol, 13% from a resin loading of 0.87 mmol/g).

**RP-HPLC:** $t_R$ = 5.1 min (RP-A).

**RP-prep.HPLC:** Prep.-A.

**$^1$H-NMR (MeOH-d$_4$, 600 MHz, 297 K)** $\delta$ = 0.78 - 1.00 (m, 36 H), 1.05 (d, $J$ = 6.6 Hz, 3 H), 1.14 (d, $J$ = 6.3 Hz, 3 H), 1.20 (m, 1 H), 1.24 - 1.39 (m, 18 H), 1.42 - 1.73 (m, 14 H), 1.80 (m, 2 H), 1.88 - 2.00 (m, 2 H), 2.05 (m, 1 H), 2.29 (m, 3 H), 2.43 (m, 1 H), 3.46 (dd, $J$ = 11.4, 7.4 Hz, 1 H), 3.62 (dd, $J$ = 11.4, 5.5 Hz, 1 H), 3.72 (dd, $J$ = 9.1, 4.5 Hz, 1 H), 3.88 (dd, $J$ = 11.6, 3.6 Hz, 1 H), 3.95 (m, 1 H), 4.04 (dd, $J$ = 11.6, 6.1 Hz, 1 H), 4.20 (t, $J$ = 10.3 Hz, 1 H), 4.34 (m, 1 H), 4.39 (m, 1 H), 4.41 (m, 1 H), 4.47 (m, 1 H), 4.67 (m, 2 H), 5.08 (d, $J$ = 9.7 Hz, 2 H), 5.41 (q, $J$ = 6.3 Hz, 1 H), 7.40 (d, $J$ = 10.3 Hz, 1 H), 7.74 (d, $J$ = 9.7 Hz, 1 H), 8.21 (d, $J$ = 9.2 Hz, 1 H), 8.62 (m, 2 H), 8.71 (d, $J$ = 7.7 Hz, 1 H), 8.81 (d, $J$ = 5.3 Hz, 1 H), 8.89 (d, $J$ = 7.9 Hz, 1 H), 9.32 (d, $J$ = 7.7 Hz, 1 H) ppm. (COOH, partial CONH and OH not assigned).

**HRMS (ESI-TOF)** calculated for $C_{64}H_{114}N_{10}O_{17}$ $[M+H]^+$ 1295.8436; found 1295.8440.

### 3. Copies of NMR spectra

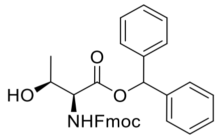


S-**1**

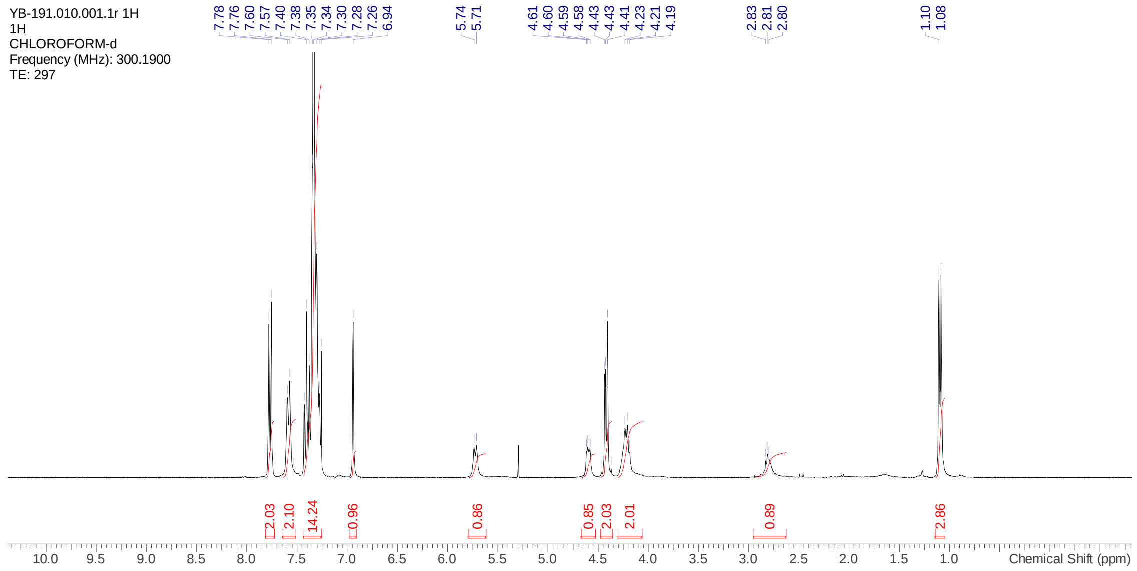


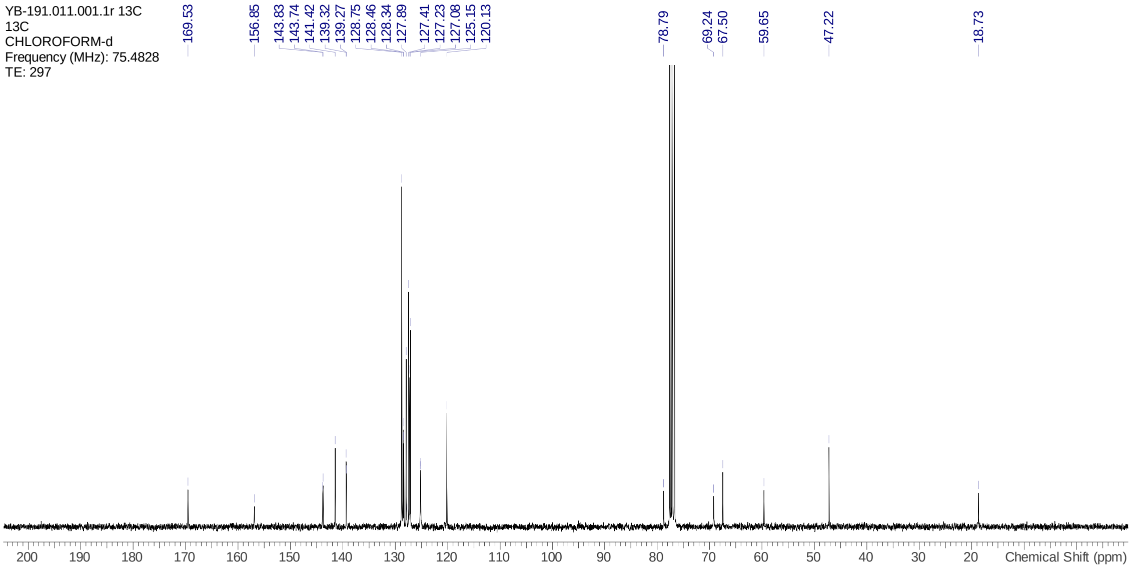

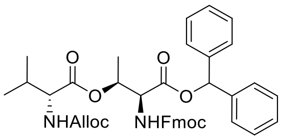
O
O
O
O
NHAlloc
NHFmoc

S-**2**

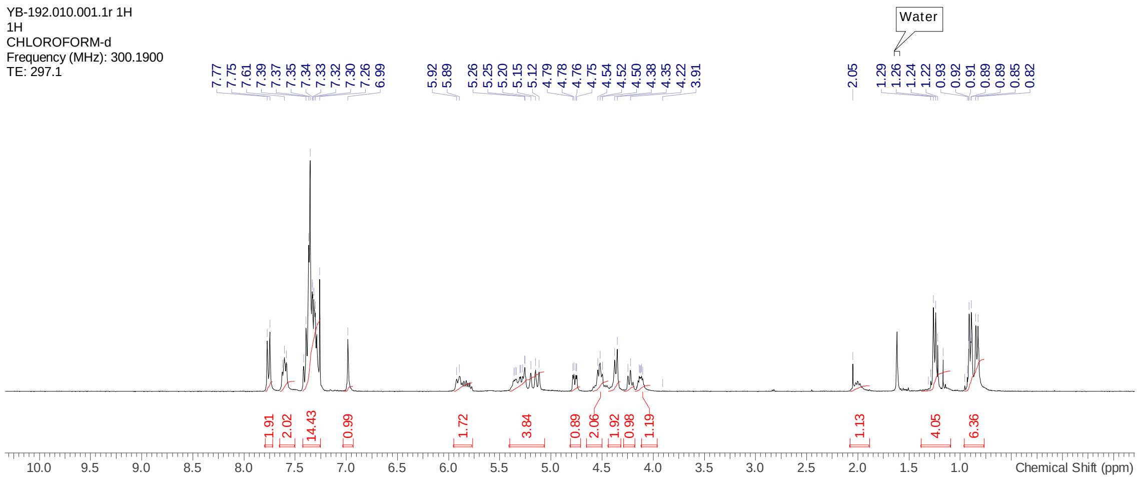
YB-192.010.001.1r 1H
1H
CHLOROFORM-d
Frequency (MHz): 300.1900
TE: 297.1
7.77 7.75 7.61 7.39 7.37 7.35 7.34 7.33 7.32 7.30 7.26 6.99
5.92 5.89
5.26 5.25 5.20 5.15 5.12 4.79 4.78 4.76 4.75 4.54 4.52 4.50 4.38 4.35 4.22 3.91
2.05
1.29 1.26 1.24 1.22 0.93 0.92 0.91 0.89 0.89 0.85 0.82
Water
1.91 2.02 14.43 0.99 1.72 3.84 0.89 2.06 1.92 0.98 1.19 1.13 4.05 6.36
Chemical Shift (ppm)

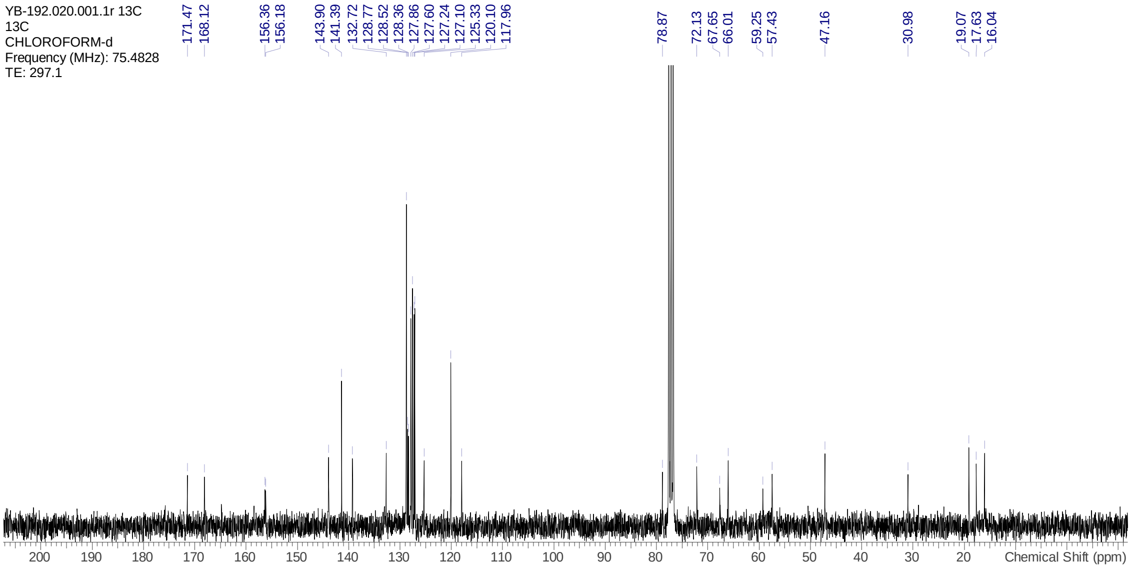
YB-192.020.001.1r 13C
13C
CHLOROFORM-d
Frequency (MHz): 75.4828
TE: 297.1
171.47 168.12
156.36 156.18
143.90 141.39 132.72 128.77 128.52 128.36 127.86 127.60 127.24 127.10 125.33 120.10 117.96
78.87
72.13 67.65 66.01
59.25 57.43
47.16
30.98
19.07 17.63 16.04
Chemical Shift (ppm)

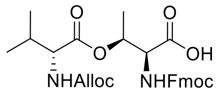
O
O
O
OH
NHAlloc
NHFmoc

S-**3**

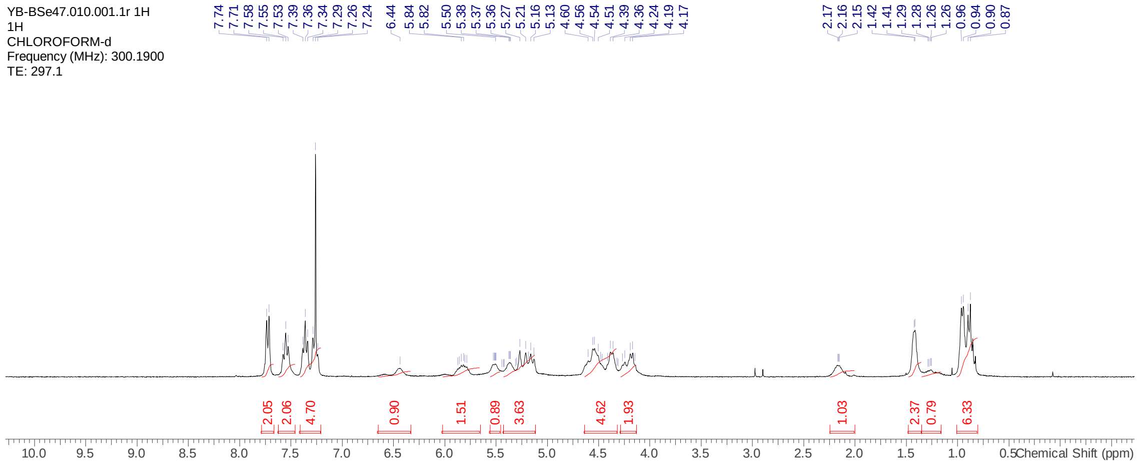
YB-BSe47.010.001.1r 1H
1H
CHLOROFORM-d
Frequency (MHz): 300.1900
TE: 297.1
7.74 7.71 7.58 7.55 7.53 7.39 7.36 7.34 7.29 7.26 7.24 6.44 5.84 5.82 5.50 5.38 5.37 5.36 5.27 5.21 5.16 5.13 4.60 4.56 4.54 4.51 4.39 4.36 4.24 4.19 4.17
2.17 2.16 2.15 1.42 1.41 1.29 1.28 1.26 1.26 0.96 0.94 0.90 0.87
2.05 2.06 4.70 0.90 1.51 0.89 3.63 4.62 1.93 1.03 2.37 0.79 6.33
Chemical Shift (ppm)

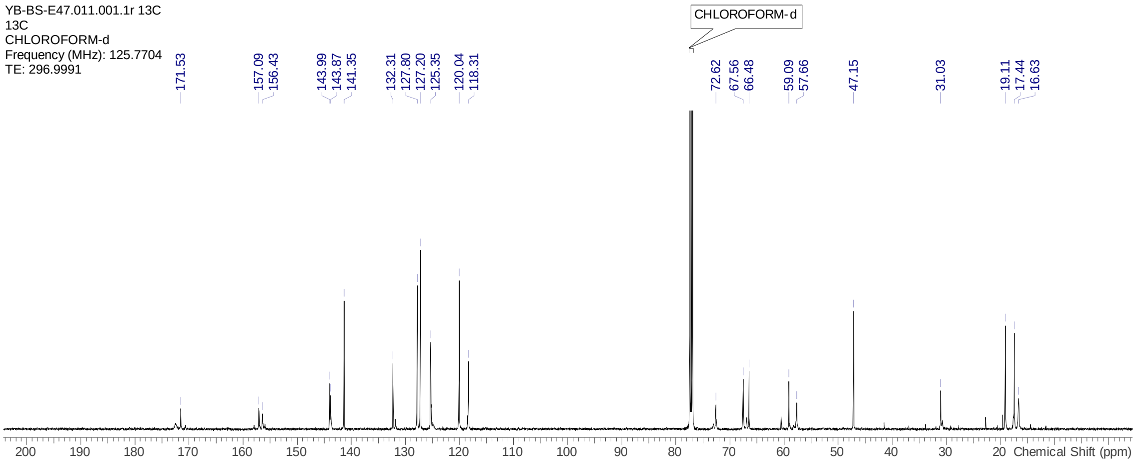
YB-BS-E47.011.001.1r 13C
13C
CHLOROFORM-d
Frequency (MHz): 125.7704
TE: 296.9991
CHLOROFORM-d
171.53 157.09 156.43 143.99 143.87 141.35 132.31 127.80 127.20 125.35 120.04 118.31 72.62 67.56 66.48 59.09 57.66 47.15 31.03 19.11 17.44 16.63
Chemical Shift (ppm)

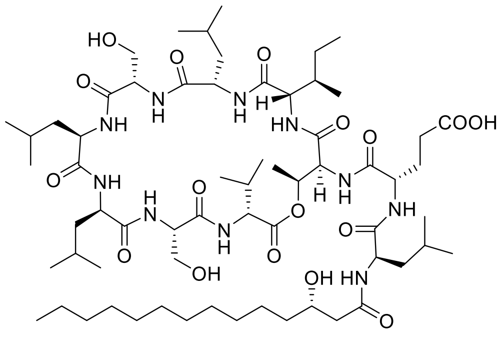


*ent*-orfamide A
(*ent*-**1**)

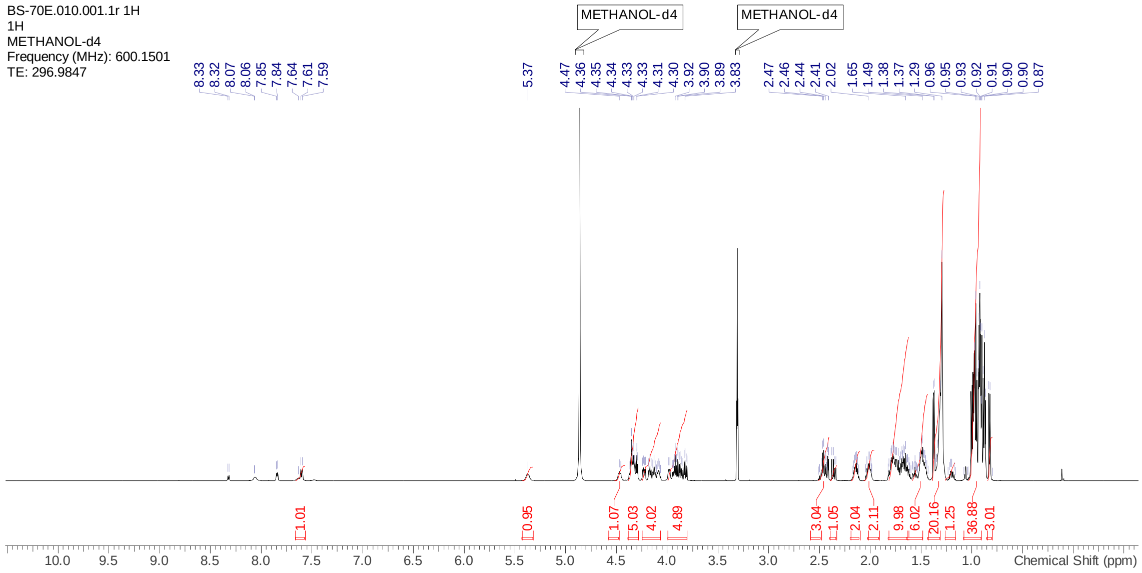


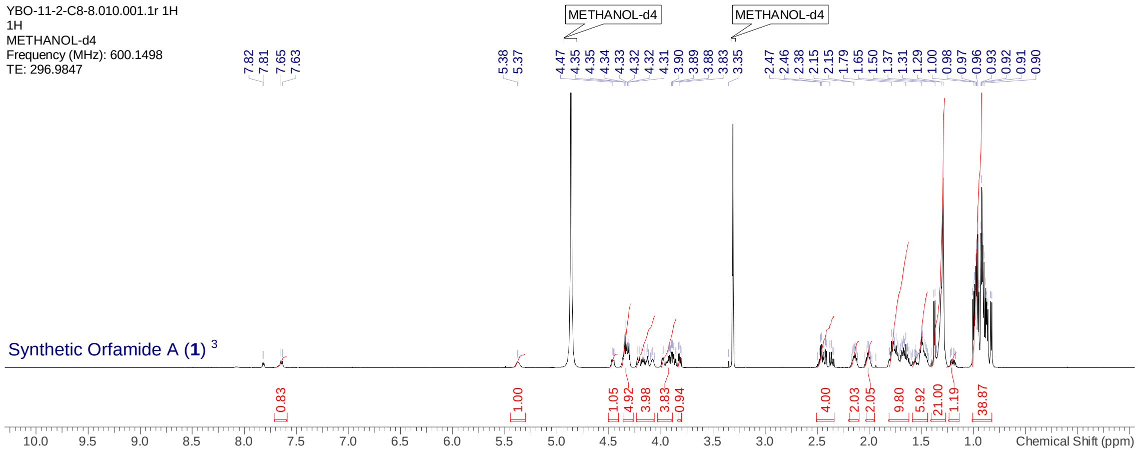

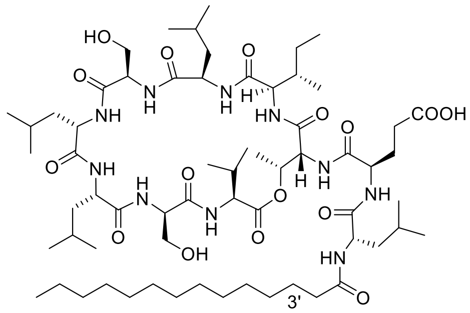


3'-deoxyorfamide A (**2**)

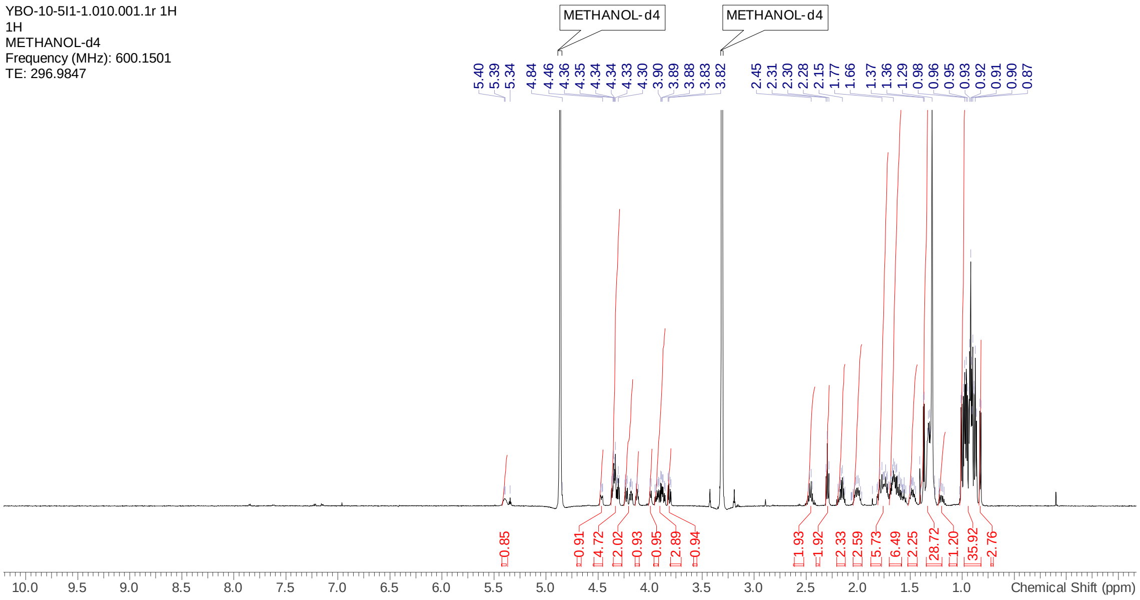

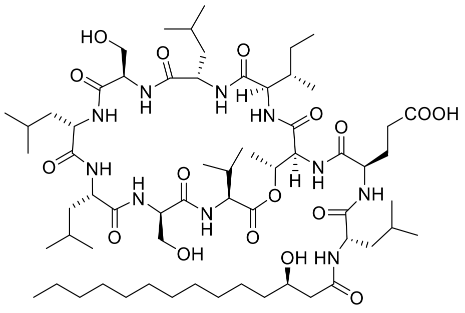


**6**

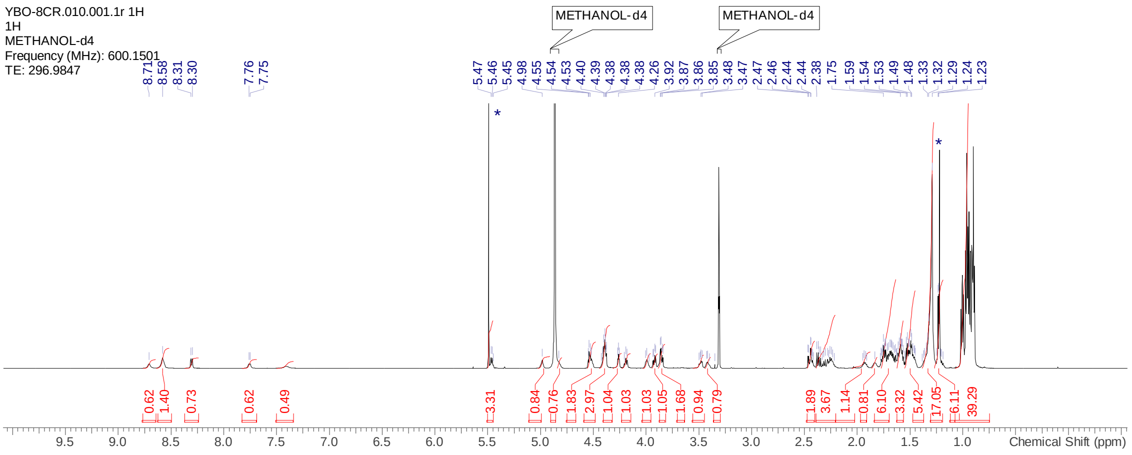


*Residual solvent peak

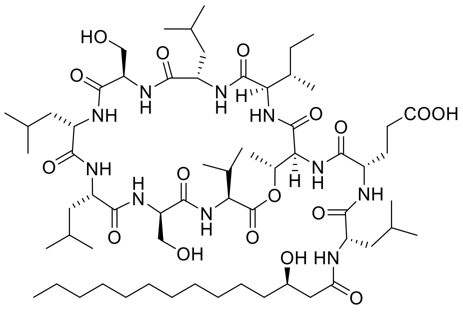

**7**

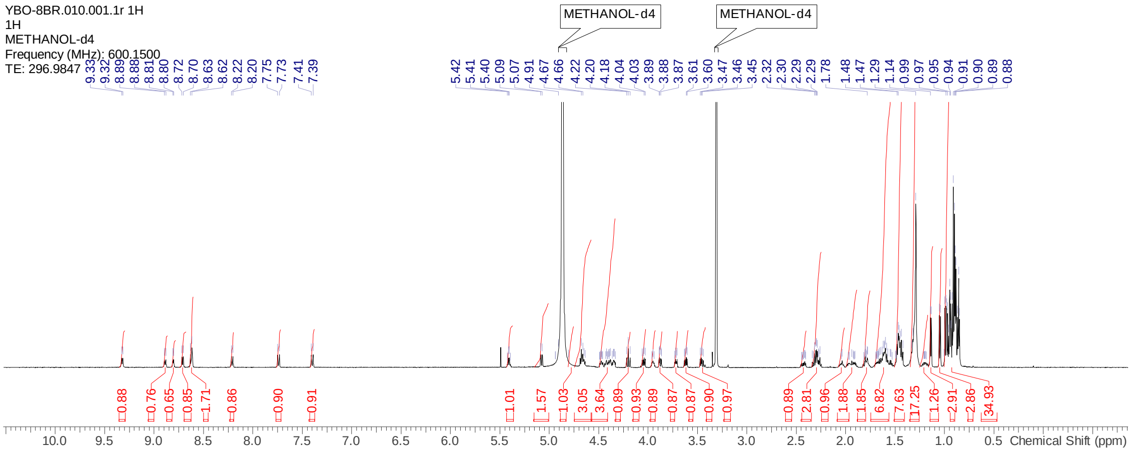

YBO-8BR.010.001.1r 1H
1H
METHANOL-d4
Frequency (MHz): 600.1500
TE: 296.9847
METHANOL-d4
METHANOL-d4
Chemical Shift (ppm)

### 4. Biological Methods

#### 4.1 $Ca^{2+}$ assay with an aequorin reporter system

The experiments were adapted from Aiyar et al., (2017).[6] Cells from the transgenic line AEQ34, expressing apo-aequorin,[6] were grown in TAP medium for 48 hours at 23 °C within a light-dark (LD) 12:12 cycle, and then incubated with coelenterazine overnight. The cell density was adjusted to 4-5 x $10^6$ cells/mL before the measurement. Obtained relative luminescence unit (RLU) were converted to molar $Ca^{2+}$ concentrations as described in Fricker et al., (1999).[7]

To measure the effect of TRP channel modulators 2-APB (Santa Cruz Biotechnology) and GsMTx4 (Smartox Biotechnology, France), the AEQ34 cells were washed twice and resuspended in HEPES buffer (10 mM HEPES, pH 7.4, 1 mM $CaCl_2$, 50 µM $MgCl_2$ and 1 mM KCl). The cells were incubated with different concentrations of modulators for 1 min before 5 min of background measurement.

#### 4.2 Deflagellation assay

The experiments were performed according to Aiyar et al., (2017)[6] with some modifications. Cells from the *C. reinhardtii* wild-type strain *C. reinhardtii* SAG 73.72 cells were treated with 5 µM of compounds for 1 min followed by fixation with 10% (v/v) Lugol's Solution (MERCK, Germany). The cells were imaged with a DIC microscope (Axiophot, Zeiss).

To measure the effect of TRP channel modulators on orfamide A-induced deflagellation, the cells were washed twice and resuspended in HEPES buffer before the experiments. The cells were incubated with TRP channel modulators for 6 min before treating with 5 µM of commercial orfamide A.

#### 4.3 Video records

Each compound was added by applying 1 µL of the compound at 110 µM to 10 µL of culture with a cell density of 4-6 x $10^6$ cells/mL, resulting in a final concentration of 10 µM which is similar to the biological concentration of orfamide A. The effects were recorded on videos without interruption. The recording started 30 sec before the addition of the compound to get a motility baseline. After 30 sec, the compound of interest was added and then 5 min were recorded to reveal potential effects. The videos were recorded in biological triplicates, with one of the replicates performed at a different day to verify reproducibility. All the videos were taken latest at midday (LD6) to ensure good motility of the cells. As positive control, commercial orfamide A was used with each video and as negative control, methanol, the solvent used to apply compounds, was used. In parallel, $Ca^{2+}$ assays were performed as described before, but running one sample at a time and using the luminometer's injector to record the whole reaction from T=0 by avoiding the delay of manually loading the plate into the machine. The taken videos were thus synchronized with the $Ca^{2+}$ measurements using Adobe After Effects CC.

#### 4.4 Phylogenetic analysis

The phylogenetic tree was built using the transmembrane (TM) domains determined by Huffer et. al (2020)[8] from all 35 structurally characterized TRP channels as a reference to identify the TM domains in the 27 TRP channels from *C. reinhardtii* (Table S2) and all the other sequences used in the phylogenetic tree (Table S3). After identifying and trimming the TM domain, a multiple sequence alignment was performed using MAFFT-DASH[9] and a phylogenetic tree was generated using IQ-TREE2[10] (Fig. 3a). Related scripts and resulting files can be found in github (https://github.com/houyu-RSC-publication/Cre_TRP_Structural_Alignment)

**Table S2**: Predicted TRP channels present in *Chlamydomonas reinhardtii* according to literature.

| Gene locus* | Protein name | Reference | Comments |
|---|---|---|---|
| Cre10.g452950 | TRP1 | [8,11–18] | |
| Cre16.g655950 | TRP2 | [11–13,15,16,18] | |
| Cre07.g327750 | TRP3 | [12,13,16–19] | |
| Cre17.g702250 | TRP4 | [12,13,18,20] | |
| Cre09.g398400 | TRP5 | [11–13,15,16,18] | |
| Cre07.g334300 | TRP6 | [12,13,15,16,18] | |
| Cre08.g358575 | TRP7 | [12,16–18] | |
| Cre03.g145147 | TRP8 | [12,13,16,18] | |
| Cre16.g658900 | TRP10 | [12,13,16,18] | |
| Cre07.g341350 | TRP11 | [11–18] | |
| Cre03.g153950 | TRP12 | [12,13,18,19] | |
| Cre03.g175050 | TRP13 | [11–13,15–18] | |
| Cre09.g397142 | TRP15 | [12,13,16,18] | |
| Cre06.g278226 | TRP16 | [11,13,15–18] | Called TRP9 in[18] and[17] |
| Cre10.g434600 | TRP21 | [13,15,16,18] | Called FAP148 in[18] |
| Cre02.g112200 | TRP22 | [15,16] | |
| Cre09.g390578 | TRP23 | [12,15,16] | |
| Cre01.g029400 | TRP24 | [16] | |
| Cre12.g529850 | TRP26 | [12,13,16,18] | Called TRP14 in[18] |
| Cre12.g493050 | TRP27 | [16] | |
| Cre10.g422750 | TRP28 | [11,13,15,16,18] | Called TRP15 in[15]. Called MOT10 in[18] |
| Cre12.g493300 | TRP29 | [12] | Named in this study |
| Cre10.g420500 | TRP30 | [12] | Named in this study |
| Cre01.g052750 | TRP31 | [12] | Named in this study |
| Cre08.g358573 | TRP32 | [13] | Named in this study |
| Cre17.g728450 | TRP33 | [12] | Named in this study |
| Cre17.g715300 | TRPP2 | [11,13,15,16,18] | |

* Gene locus according to the *C. reinhardtii* genome v5.6 from the Joint Genome Institute (JGI)
